# MUC2 Expression Modulates Immune Infiltration in Colorectal Cancer

**DOI:** 10.1101/2024.08.06.594842

**Authors:** Christophe M. Raynaud, Ayesha Jabeen, Eiman I. Ahmed, Satanay Hubrack, Apryl Sanchez, Shimaa Sherif, Ahmad A Al-Shaibi, Jessica Roelands, Bernice Lo, Davide Bedognetti, Wouter Hendrickx

## Abstract

**Introduction:** Colorectal cancer (CRC) is a prevalent malignancy with significant morbidity and mortality worldwide. A deeper understanding of the interaction of cancer cells with other cells in the tumor microenvironment is crucial for devising effective therapeutic strategies. MUC2, a major component of the protective mucus layer in the gastrointestinal tract, has been implicated in CRC progression and immune response regulation.

**Method:** In this study, we sought to elucidate the relationship between MUC2 expression and immune infiltration within CRC, using *in-vitro* models involving two well-established cell lines, HT-29 and LS-174T. By employing CRISPR-mediated MUC2 knockout, we investigated the influence of MUC2 on tumor immune infiltration and its interplay with T cells and NK cells enriched peripheral blood mononuclear cells (PBMCs) in 3D spheroid cultures.

**Results:** While MUC2 was more abundant in LS-174T cell lines compared to HT-29, its knockout resulted in increased immune infiltration solely in the HT-29 cell line, but not in LS-174T. We revealed that the removal of MUC2 protein was compensated in LS-174T by the expression of other gel forming mucin proteins (Muc6, Muc5B) commonly expressed in gastrointestinal epithelium, while this was not observed in HT-29 cell line.

**Discussion:** We propose that the role of MUC2 documented in CRC progression can partially be explained by impairing immune infiltration due to physical barrier established by the gel forming proteins such as MUC2 in mucinous CRC. On the other hand, the removal of MUC2 expression can be compensated by alternative gel forming mucin proteins, thereby impeding any increase in tumor immune infiltration.

## Introduction

Colorectal cancer (CRC) is a complex disease influenced by various genetic, epigenetic, and environmental factors^1–3^. While significant advancements have been made in understanding CRC biology, the role of the tumor microenvironment and its interactions with tumor cells remains an area of active investigation. The tumor microenvironment plays a critical role in modulating tumor growth, invasion, and immune surveillance, ultimately determining the overall clinical outcome of CRC patients^1^.

Mucin plays an important role in the colon as it is the primary component of the protective mucus layer lining the colon’s surface. This mucus layer acts as a barrier against harmful substances, bacteria, and pathogens, preventing them from directly interacting with the colon’s epithelial cells^4^. Additionally, mucin aids in lubricating and facilitating proper bowel function. MUC2 is an essential member of the mucin family in colon as it is a gel forming mucin along with MUC5AC, MUC5B, MUC6 and MC19^5^. Aberrations in MUC2 expression have been implicated in various gastrointestinal disorders and malignancies, including colorectal cancer (CRC)^6^. Notably, changes in MUC2 expression have been associated with CRC progression and prognosis, where patients with low-expressed MUC2 CRC tumor are significantly linked with poor overall survival highlighting its potential significance as a prognostic biomarker^7^. Nevertheless, the impact of MUC2, the aberrant glycosylation of mucins and its interaction with immune system in CRC is an active area of investigation.

In the context of CRC subtypes, mucinous CRCs are characterized by elevated MUC2 expression compared to normal colon, serving as a hallmark feature^8–10^. Conversely, non-mucinous CRCs typically exhibit decreased MUC2 expression^8^. The loss of MUC2 expression has been linked to increased proliferation of intestinal epithelial cells in response to mucosal inflammation^11^.

Multiple factors regulate MUC2 expression in colonic epithelial cells. Studies have shown that methylation of the MUC2 gene promoter is significantly lower in mucinous CRC lines compared to non-mucinous lines, correlating with mucin protein expression^12^. The tumor suppressor protein p53 is another crucial regulator of MUC2 expression, with loss of functional p53 leading to downregulation of MUC2^13^. This transcriptional regulation of MUC2 by p53 has been observed in various cell lines^14^, consistent with the reduced incidence of p53 mutations in mucinous carcinomas^13^. On the other hand, non-mucinous CRCs often display high rates of p53 mutation and low MUC2 expression^14^. Moreover, the mitogen-activated protein kinase (MAPK) signaling pathway has been identified as another regulator of MUC2 gene transcription^15^. Notably, MAPK signaling is upregulated in CRCs harboring KRAS mutations and in the context of chronic inflammation, both of which are common features of mucinous cancer^16^.

In summary, MUC2’s critical role in the gastrointestinal tract, its association with CRC progression, and its regulation by various factors underscore the importance of investigating its impact on the immune landscape within the tumor microenvironment. We previously found that, overall, mucinous colon tumors are characterized by a reduced Th1-oriented immune response transcriptomically determined by the Immunologic Constant of Rejection (ICR)^1^. Considering the described barrier function of the mucus layer, we proposed that infiltration of immune cells to the tumor might be hampered in mucinous cancers. Therefore, understanding these intricate relationships may pave the way for novel therapeutic approaches targeting MUC2 and its interplay with the immune system to improve outcomes for CRC patients.

In this study, we aimed to investigate the role of MUC2 in CRC immune infiltration, focusing on the interaction between MUC2-altered CRC cells and T cells and NK cells. We utilized CRISPR-mediated gene editing techniques to specifically knockout MUC2 expression in HT-29 and LS-174T cell lines, two widely studied CRC models known for their distinct characteristics. These cell lines provided an ideal platform to assess the impact of MUC2 modulation on tumor infiltration and its interplay with immune cells in a controlled and reproducible in vitro setting. Additionally, to better recapitulate the tumor microenvironment, we employed 3D spheroid cultures, a well-established model that more closely mimics the cellular architecture and interactions present in solid tumors than a traditional monolayer. By appending T cells and NK cells enriched PBMCs into the 3D spheroid cultures, we aimed to simulate the immune-rich environment encountered in the CRC microenvironment.

## Material and methods

### Cell lines and culture

LS-174T (CL-188), HT-29 (HTB-38) and LoVo (CCL-229) cell lines were purchased from ATCC. Original and modified cell lines were cultured in advanced RPMI (Gibco, #12633012) complemented with 10% FBS (Sigma, # F4135-500ML), Glutamax (Gibco, #35050061) and antibiotic-antimycotic (Gibco, #15240096) (complete media). Cells were cultured at 37°C, 5% CO2 and 95% humidity. Cells were detached using TrypLE express enzyme (Gibco, #12605036). For 2D culture, T-75 (Falcon, #353136) or 96 well plates (Falcon, #353072) were used. For 3D culture low adherence NunclonSphera U bottom 96 well plates were used (Thermo, #174925).

### RNA extraction

After culture and detachment cells were rinsed in DPBS (Gibco, #10010-031) dry cell pellet was then used for RNA extraction as follow. RNA was extracted Maxwell® RSC simplyRNA Cells Kit (Promega, #AS1390) according to manufacturer recommendations using Maxwell® RSC Instrument (Promega, #AS4500). RNA was recovered with 50ul of RNase free water and measured with QuantiFluor® RNA System (Promega, #E3310). RNA was stored at −80°C until use.

### Q-PCR

1ug of RNA was used for reverse transcription using TaqMan™ Reverse Transcription Reagents (Invitrogen, #N8080234) using random hexamer following manufacturer’s recommendation. cDNA was diluted 20 times with DNA/RNA free water. TaqMan™ Gene Expression Master Mix (Applied bioscience, #4369016) was used together with Hs03003631_g1 (for Eukaryotic 18S rRNA) and Hs03005103_g1 (for MUC2) TaqMan probes (Thermo scientific, #4331182). According to manufacturer recommendation. Real time PCR was run in 96 well plates on QuantStudio 12K flex system (Thermofisher Scientific). Q-PCR was done in triplicate for each sample and data were analyzed by gene expression comparison using ΔΔCT on (QuantStudio 12K Flex Realtime PCR system V1.2.2).

### Western blot

After culture, 5 × 10^6^ cells of cells for 2D culture or 30 spheroids at day 5 for 3D culture were harvested and lysed in lysis buffer containing (10 mmol/L HEPES, 150 mmol/L NaCl, 1 mmol/L ethylene glycol-bis(β-aminoethyl ether)-*N,N,N′,N′*-tetraacetic acid, 0.1 mmol/L MgCl_2_, 0.5% Triton X-100, pH 7.4) containing 1 mM PMSF and 10 mM N-ethyl maleimide (NEM). 50 mg lysate as input for each condition were loaded onto 4%–15% TGX gels (Bio-Rad, Hercules, CA). Proteins then were transferred onto nitrocellulose membrane and blotted with antibodies against proteins of interest. MUC2 antibody (CCP58) (Novus #NBP2-25221) and HRP-conjugated β actin monoclonal antibody (Proteintech, #HRP-60008) were used as primary antibody as well as Anti-mouse IgG (H+L) antibody, human serum adsorbed and peroxidase-labeled (Seracare, # 5450-0011) as secondary anti body.

### CRISPR cell engineering

#### gRNA design

MUC2 gene sequence was obtained from ensembl human genome assembly GRCh38.p14 HSCHR11_3_CTG1: 149,268-178,812. IDT custom gRNA design tool was used to design gRNA along exon 2 of transcript ID (ENST00000643422.1).

5 gRNA were designed as follow:

**Table 1:**
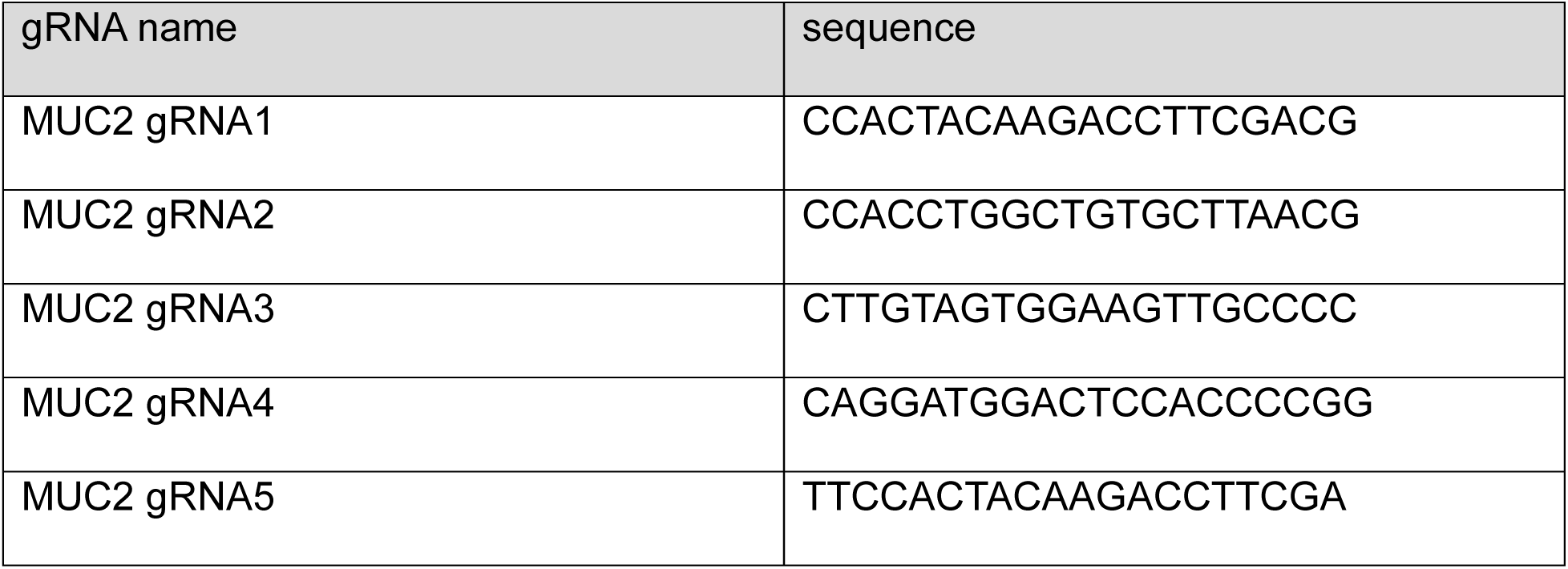
sequence of each gRNA designed for MUC2 K.O.

The location along the exon2 of MUC2 of each gRNA is shown in supplementary figure 1A. Forward and reverse primers were ordered accordingly to be inserted in the appropriate plasmids as described below.

#### In-Vitro gRNA synthesis

The gRNAs were synthesized using the GeneArt™ Precision gRNA Synthesis Kit as per manufacturer recommendation. The concentration of gRNA was determined by the Qubit® RNA BR Assay Kit as per manufacturer recommendation and diluted at 250ng/ul final concentration and stored at −80C until use [3-5].

#### Cell transfection

Electroporation was performed using Neon transfection system (Invitrogen, #MPK5000) with Neon™ Transfection System 10 µL Kit (Invitrogen, #MPK1096) using 1ug of DNA for 1.10^5^ cells in 24 well plate according to the manufacturers recommendation. Electroporation protocol for each cell line was identified using pmaxCloningTM vector (Lonza, # VDC-1040). The optimal protocol for each cell line is indicated in table 2.

**Table 2:** optimal electroporation setup for each cell line.

| CELL LINE | VOLTAGE (V) | WIDTH (ms) | PULSES |
| --- | --- | --- | --- |
| HT-29 | 1700 | 20 | 1 |
| LS-174T | 1400 | 20 | 2 |

#### Bulk analysis

72h after electroporation, half of the cells were collected and washed with DPBS and then lysed in 10ul cell lysis buffer complemented with 0.4ul of protein degrader, from GeneArt™ Genomic Cleavage Detection Kit (Thermofisher scientific, #A24372). PCR was performed directly on cell lysis using the primers provided in table 3 and relative location of primers to gRNA is provided in Supplementary Figure 1A, with Amplitaq gold 360 master mix (applied biosystem, #4396790) as per manufacturer recommendation with annealing at 60C and elongation step of 30 seconds.

**Table 3:**
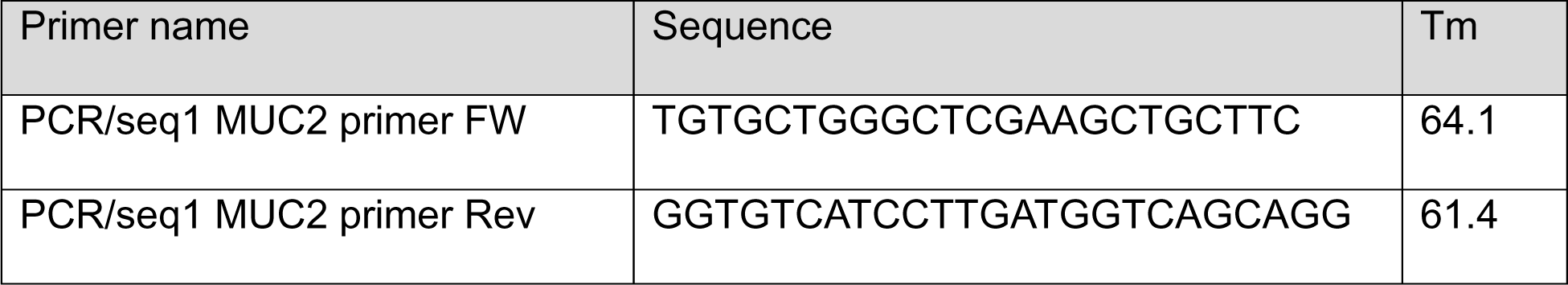
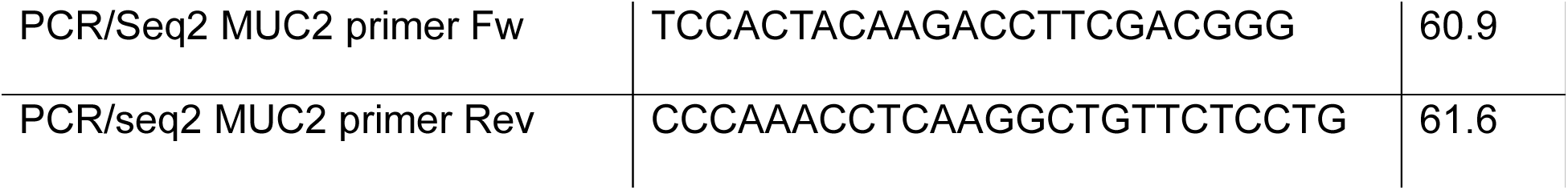
primers used for PCR and sanger sequencing of MUC2 exon 2.

The PCR product was used for sanger sequencing and analysis as described below.

#### Sanger sequencing

The same primers used for PCR were used for sanger sequencing. Briefly, PCR product were purified using Exosap-IT (Thermofisher scientific, # 75001) followed by labeling using BigDye™ Terminator v3.1 Cycle Sequencing Kit (Thermofisher, # 4337455) as per manufacturer recommendation. Sanger products were cleaned using DyeEx™ 96 kit (Qiagen, #63204) as per manufacturer recommendation. Sanger sequences were recorded on ABI3500 sequencer (Applied biosystem, #4406016).

#### Sanger sequence analysis

Synthego ICE https://ice.synthego.com/#/ [6] online tool was used to analyze the indels in bulk and for each individual clones (after sort described below). The selected clone had indel on both allele indel non multiple of 3 inducing frame shift and therefore K.O.

#### Single cell sorting

Cells were harvested and blocked in PBS with 5%FBS and 1%BSA and cell clumps removed on 40uM cell strainer (Falcon, #382235). Single-cell suspension was analyzed and sorted on SORP FACSAriaIII (BD Biosciences Special Order Research Product). Data were processed with BD FACSDiva™ Software V8.0.1 (BD Biosciences). Doublets were excluded by FSC-W × FSC-H and SSC-W × SSC-H analysis. A dumping channel was used to eliminate auto-fluorescent cells with 405 nm violet laser and 525/50 emission filter. During cell-sorting single-cell sort mask was applied and 1 cell was seeded per well of 96 well plate in 200µl of compete media supplemented with 10µM of rock inhibitor (Y-27632) (Stem cell technologies, #72305).

Each clone was further amplified sequentially in 96, 48, 24, 6 well plate and T25, T75 flask before being analyzed individually to assess the genomic edition by sanger sequencing after DNA extraction.

#### DNA isolation of clones

DNA isolation was performed using Maxwell® RSC Cultured Cells DNA Kit (Promega, # AS1620) according to manufacturer recommendations using Maxwell® RSC Instrument (Promega, #AS4500). DNA was recovered with 100ul of elution buffer and measured with QuantiFluor® ONE dsDNA System (Promega, # E4871). DNA was stored at −80°C until use.

#### GFP expression in control and K.O. cell lines

To help with growth analysis, co-culture, video-microscopy, and viability measurement cancer cells (both control and MUC2 K.O. cell lines) were further modified to express GFP under CMV promoter. The modification of cells was done using Cyagen EGFP lentivirus (Cyagen, #LV-EGFP-0102, lot#140224LVT02) as per manufacturer recommendations. After culture, cells expressing GFP were further purified by cell sorting in bulk as previously described where GFP fluorescence was acquired with 488 nm blue laser and 530/30 nm emission filter. During cell-sorting 4-way sort mask was applied. To ensure maximum purity, cells were serially sorted 3 times prior analysis and use.

#### WES

Capillary western blot (WES) was performed for the analysis of E-Cadherin, N-cadherin and vimentin, to see the potential EMT modification of the MUC2 K.O. cells. After culture, 5 × 10^6^ cells were washed with DPBS and lysed with 400ul of RIPA Lysis and Extraction Buffer (Thermo Scientific, #89900) complemented with Halt™ Protease Inhibitor Cocktail (Thermo scientific, #78430) and sonication for 30 sec. cell debris was removed by 30 min centrifugation at 14.000g. Supernatants containing protein extract were kept at −20°C until use. Protein concentration was assessed using Pierce BCA protein assay kit (Thermo scientific, #23225). Capillary western blot was done using a Wes system (protein simple) with 12-230 kDa Separation Module, 8 x 25 capillary cartridges (Protein simple, #SW-W004), EZ Standard Pack 2 (Protein simple, #PS-ST02EZ-8) and Anti-mouse detection module (Protein simple DM-002). Mouse anti human E-Cadherin (R&D systems, #MAB1838) diluted at 0.5ug/ml, mouse anti human N-Cadherin (Novus Biologicals, #NBP1-48309) diluted at 1 in 10, mouse anti human Vimentin (Novus Biologicas, #NBP1-92687) diluted at 1 in 25 and anti β-actin (Licor, #926-42212) diluted at 1 in 100 were used as primary antibody.

Analysis was done using compass for Simple western (ProteinSimple, V5.0.0) and area of histogram peaks were used for quantification. All western blot analysis were normalized for β-actin expression.

### PBMCs isolation, enrichment, activation, and staining

#### PBMCs isolation

Whole blood from healthy donor was collected in Sidra medicine under the IRB 1500815-2 was collected in EDTA tubes (XXX) and immediately processed for PBMCs isolation. Blood was diluted to 1 :1 with DPBS (Gibco, #10010-031) and gradient dilution was performed with SepMate™-50 (IVD) tubes (stem cells technologies, # 85450) containing 15ml of Lymphoprep (stemcell technologies, # 07801), according to manufacturer recommendations. Briefly cells were centrifuged in sepmate tube for 10 min at 1200g. Supernatant and PBMCs were collected in a fresh tube. PBMCs were then rinsed twice with DPBS by centrifugation 8 min at 300g. PBMCs were then used for magnetic beads enrichment of T and NK cells as described below.

#### PBMCs enrichment

PBMCs isolated as previously described were further enriched in T cells and NK cells by elimination of CD14 and CD19 positive cells using MACS cell separation according to manufacturer recommendations. Briefly, cells in 80ul buffer per 10^7^ cells and 20ul of CD14 microbeads (Miltenyi, #130-050-201) and CD19 microbeads (Miltenyi, #130-050-301) were incubated for 15 min at 4C. Cells were washed with 2ml of buffer and centrifuged for 8min at 300g. The positive fraction was removed by separation on LS columns (Miltenyi, # 130-042-401) on QuadroMACS separator (Miltenyi, # 130-091-051) with 3 washes with 3ml of buffer. Non-labeled cells were then pelleted and stored at −80°C in 90% FBS + 10% DMSO until use.

#### T cell activation

T cells in the enriched fraction of PBMCs were thawed and activated prior to co-culture, in 2D or 3D, with cancer cells using Dynabeads™ Human T-Activator CD3/CD28 (Gibco, #1132D) as recommended by the manufacturer. Enriched and activated PBMCs are referred to as E/A-PBMCs.

#### T cell staining for video microscopy

When performing live imaging, enriched/activated PBMCs were stained with CellTracker™ Red CMTPX Dye (thermofisher scientific, #C34552) according to manufacturer recommendations prior to co-culture.

### 2D Co-culture protocol and analysis

Co-culture in 2D was performed as sketched in Supplementary Figure 2A. A total of 500 GFP cancer cells (HT-29 and LS-174T control (CTRL) or MUC2 K.O.) were seeded per well in a 96-well plate and cultured for 48h in 200ul of complete media. After removing 100ul per well, various numbers ranging from 500 to 10.000 activated/enriched PBMCs were re-suspended in 100ul of fresh complete media, then seeded on top of the GFP cancer cells. GFP was assessed by fluorescence intensity measurement by well scan from bottom with excitation at 494 nm and emission at 517 nm on Ensight plate reader (Perkinelmer, #HH34000000).

### 3D co-culture protocol

Co-culture in 3D was performed as sketched in Supplementary Figure 2B.2000 cancer cells per well (CTRL or K.O.) were seeded in a low adherence Nunclon Sphera 96U well plate (Thermofisher scientific, #174925) and allowed to form spheroids for 120 hours (5 days) in 200ul complete media. On day 5, 100 ul of media was removed (without disturbing the spheroids) and 4000 enriched/activated PBMCs were added in each well. GFP expression was monitored using GFP cancer cells on Ensight plate reader as previously described. Alternatively, GFP cancer cells and enriched/activated PBMCs stained with CellTracker™ Red were used for imaging over 48h on CellDiscoverer 7 (CD7, Zeiss) equipped with incubation chamber set at 37°C and 90% humidity. For flow cytometry analysis, IN and OUT fractions were separated and stained and analyzed as described below.

### IN and OUT fraction isolation

OUT and IN fraction were separated after 2 days co-culture. OUT and IN compartments were isolated by first pooling the 30 cocultures wells in FACS tubes (falcon, #). Spheroids were gently resuspended and left to sediment to the bottom of the tube. Supernatant cell suspension constituted the non-infiltrating immune cells (=OUT). These steps were repeated 2 times with PBS to wash the remaining spheroids (=IN) from the non-infiltrating immune cells. IN fraction was imaged with CD7 at this stage. Alternatively, both IN and OUT fractions were then trypsinized for 30 min to obtain a single cell suspension and both fractions were strained on 40um strainer (falcon, #) and further analyzed by flow cytometry.

### Flow cytometry analysis

Previously isolated single cell suspension from IN and OUT fraction were stained for flow cytometry analysis. LIVE/DEAD® Fixable Aqua stain (Thermofisher, # L34957) was used as per manufacturer recommendation. Then, unspecific sites were saturated by incubation for 30 min with blocking media consisting of DPBS competed with 5% FBS and 2% Bovine Serum Albumin. After removing the blocking media, spheroids were incubated 45 min with primary antibodies indicated in Table 4 diluted in blocking media.

**Table 4:** antibodies used for flow cytometry and flow cytometer analysis setup.

| Antibody | Clone | Manufacturer | Catalogue number | Laser wavelength | Band pass of analysis |
| --- | --- | --- | --- | --- | --- |
| Epcam-AF488 | 323/A3 | Invitrogen | MA5-38713 | 488 | 530/30 |
| CD45-PE-Texas Red | HI30 | Invitrogen | MHCD4517 | 488 | 615/24 |
| CD56-PE-Cy5 | 1B7 | Invitrogen | 15-0567-42 | 488 | 675/30 |
| CD4-SB600 | SK3 | Invitrogen | 63-0047-42 | 405 | 615/24 |
| CD8a-SB436 | SK1 | Invitrogen | 62-0087-42 | 405 | 445/45 |
| Live/dead (fixable aqua D) |  | Invitrogen | L34966 | 405 | 530/30 |

After staining, cells were rinsed 2 times with staining media and resuspended in a final volume of 100ul. 80ul were acquired on Acea Novocyte (Acea, # 2010050) on low speed and analyzed with Novoexpress software (Acea). Compensations were calculated using UltraComp eBeads™ Compensation Beads (Thermofisher scientific, #01-2222-42). Fluorescence minus one (FMO) were used to define the positive/negative cut off for the gating strategy presented in supplementary figure 3.

### OCT embedding and cryosection

8-12 spheroids of specified age and culture method were pulled together in 1 well of sphere 96U well plate. Media was removed completely trying not to disturb the spheroids. OCT was then poured over the spheroids and let solidify at −20°C. Section of 10µm were cut using a cryostat (Leica biosystem, #CM3050S), and section were attached onto Superfrost^TM^ plus gold slides (Epredia, #FT4981IGLPLUS-001) and stored at −20°C until staining.

### Immuno-fluorescence staining and confocal imaging

Slides were allowed to warm up at room temperature for 15 min prior staining. Area containing spheroids were surrounded with PAP pen (Sigma, # Z377821), and unspecific sites were saturated by incubation for 30 min with blocking media consisting of DPBS competed with 5% FBS and 2% Bovin Serum Albumin. After removing the blocking media, spheroids were incubated 45 min with primary antibody diluted in blocking media (mouse anti human MUC2 antibody (CCP58) (Novus #NBP2-25221) diluted at 1:100 or rabbit anti human Muc5B (Thermofisher scientific, # PA5-82342) diluted at 1:25 or Mouse anti human Muc5AC (Thermofisher scientific, # MA5-12178)). After removal of primary antibody and two rinse of the spheroids with blocking media, a 30 min with secondary antibody diluted at 1:200 in blocking media was performed (Alexa Fluor 488 (AF-488) Goat anti Mouse IgG (H&L) antibody (Thermofisher scientific, #A-11001) or AF-488 Goat anti Mouse IgG1 (Thermofisher scientific, #xxx). Finally, after two rinses with blocking media cover-slides were mounted with SlowFade™ Gold Antifade Mountant with DAPI (Thermofisher scientific, #S36942).

**Table 5:**
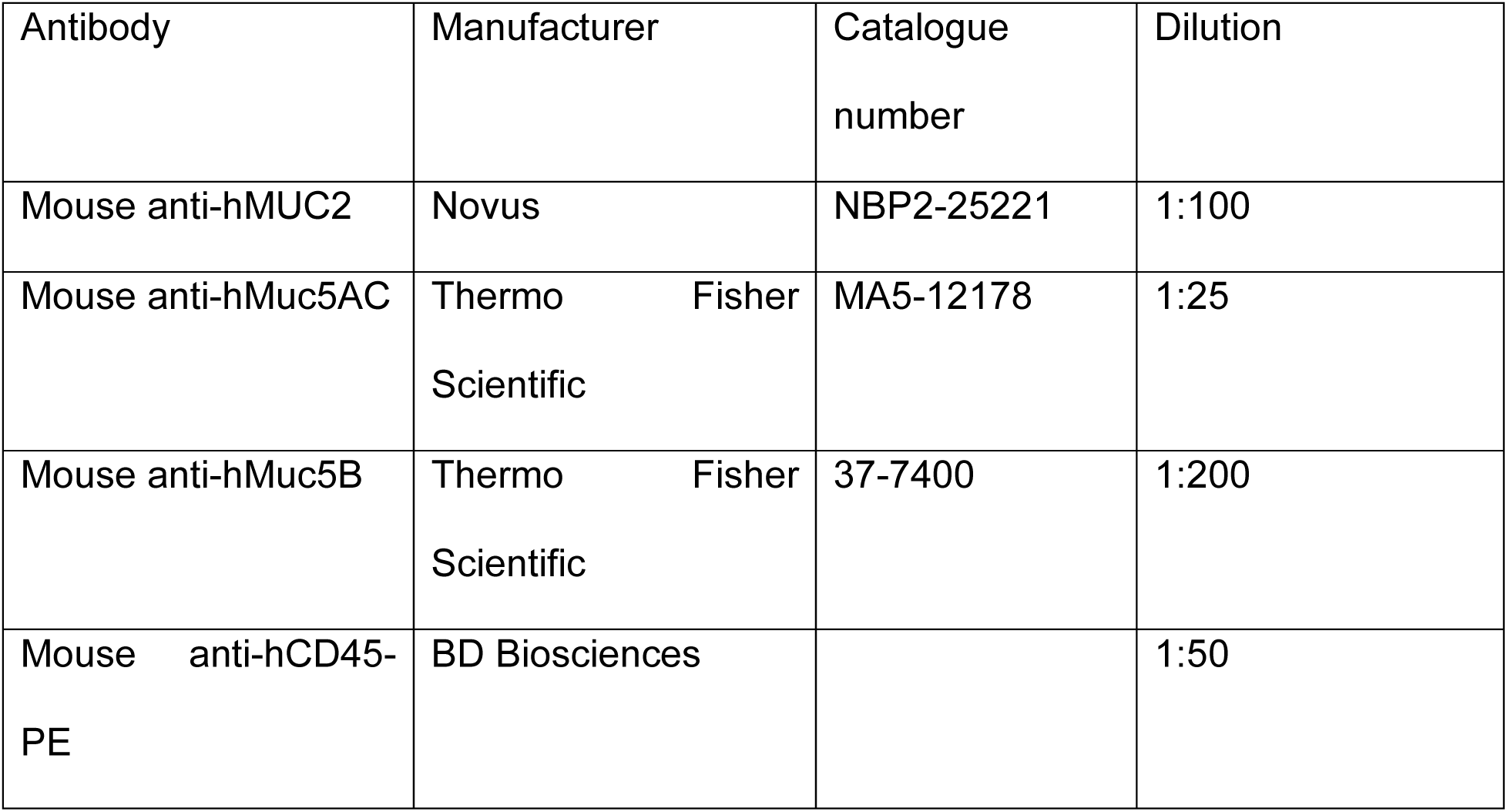
antibodies used for immunofluorescence.

Immunofluorescent stained spheroids were imaged on Zeiss confocal Microscope LSM780 (Zeiss) using 405 nm laser for DAPI analysis, intune laser at 490nm for AF-488. Z-stack of 5 were acquired and maximum intensity projection are represented.

### CD7 imaging

To study the effect of E/A PBMC’s on HT-29 and LS-174T CTRL GFP and MUC2 K.O. GFP cells were imaged for 48 hours using Zeiss Cell discoverer 7 (CD7) imaging system. Where, E/A PBMC’s were stained with cell tracker red CMTPX dye (Thermofisher scientific, # C34552) following manufacturer’s instructions. Axiocam 506 with objective lens magnification of 5X was used and detection wavelengths were 470nm for EGFP, 570nm for AsRe2; and oblique for brightfield. Image acquisition to quantify the roundness and growth of both HT-29 and LS-174T CTRL GFP and MUC2 K.O. GFP cells were also performed using CD7 with same optical settings and laser detectors. Roundness of both HT-29 and LS-174T CTRL GFP and MUC2 K.O. GFP cells were quantified using Zen Blue software to measure area (µm^2^) and perimeter (µm). Formula used to calculate roundness of a spheroid is 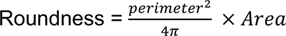.

### Imaris analysis

Spheroids growth and post rinse analysis of HT-29 and LS-174T CTRL GFP and MUC2 K.O. GFP were analyzed by measuring its area (µm^2^) using Imaris V 9.0.1 software. EGFP channel was used as a source to analyze the spheroids for both the cell lines and same parameters were used for the CTRL and MUC2 K.O.

### Plate reader analysis

GFP expression was analyzed on plates for both 2D and 3D culture setup using EnSight Multimode Plate Reader (Perkinelmer, #HH34000000). GFP was acquired by bottom excitation at 485nm and emission analysis at 519nm wavelength. A well scan was performed of 10×10 in round shape area with distance of 0.75 between measurement for 2D and 3X3 in round shape area with distance of 0.2 between measurement for 3D (centered around the U bottom well). Average intensity was measured for each well.

### RNA seq analysis

#### mRNA sequencing

Library construction and sequencing was performed at the Sidra Clinical Genomics Laboratory Sequencing Facility. Sample integrity and concentration of the Total RNA was controlled and measured using the Standard sensitivity RNA assay on the Perkin Elmer Caliper Labchip GXII. 500ng of Total RNA was used for library preparation using the Illumina Truseq Stranded mRNA kits. To obtain mRNA libraries, poly-A RNA selection was performed using an Oligo-dT magnetic bead system, followed by fragmentation, first strand synthesis using Superscript IV and second strand synthesis. The cDNA obtained after reverse transcription is then ligated with IDT for Illumina UD Indexes and amplified for 10 cycles. Library quality and concentration were then assessed using the DNA 1k assay on a Perkin Elmer GX2. Library quality and concentration were then assessed using the LabChip High Sensitivity assay on a Perkin Elmer GX2 and by qPCR using the KAPA Library quantification kit on a Roche LightCycler 480 II. The RNA libraries were sequenced with paired end 150bp on Novaseq 6000 system (Illumina, USA) following the manufacturer’s recommended protocol at depth of 40 Million Reads per sample.

Single samples were sequenced across multiple lanes, and the resulting FASTQ files were merged by sample. All samples passed FastQC (v. 0.11.8) were aligned to the reference genome GRChg38 using STAR (v. 2.6.1d) ^17^. BAM files were converted to a raw counts expression matrix using HTSeq-count (v. 0.9.1) ^18^.)

#### Data processing and normalization

Quality control (QC) check was performed using FastQC module (Python v.2.7.1, FastQC v.0.11.2) in the raw data. Adaptor sequencing trimming was run using flexbar (v.3.0.3) using Illumina primers FASTA file. Then, alignment of the reads to human reference genome GRCh38.93 was performed via Hisat2 (v.2.1.0) using SAMtools (v.1.3). QC was performed to confirm alignment quality and paired-end mapping overlap (Bowtie2, v.2.3.4.2). Finally, read counts for each gene was created using featureCounts function of subreads (v.1.5.1). Gene count was normalized using R package EDASeq (Exploratory Data Analysis and Normalization for RNA-seq) (v. 2.34.0) to correct for within lanes effect (GC content) and between lanes effect (sequencing depth). Then quantile normalization was performed on the resulting values using R package preprocessCore (v.1.62.1) and then log2 transformed. All downstream analysis was done using R programming software (v. 4.3.1, or later). Principal component analysis (PCA) was performed to evaluate global changes between samples based on gene expression using “prcomp” function and data was plotted using R CRAN package ggplot2 (v. 3.4.2). For data visualization ggplot2 (v. 3.4.2) and ComplexHeatmap (v.2.16.0) were used for boxplot and heatmap plots respectively.

#### Differentially expressed genes (DEG)

DEG analysis between groups was performed using log2 normalized expression matrix using R Bioconductor package limma (v. 3.56.2) (Ritchie et al. 2015) with Benjamini-Hochberg (B-H) FDR correction. Within each comparison, genes with row sum equal to zero were excluded. Overlapping DEGs between groups were visualized using R CRAN package Venn (v. 1.11) or volcano plot using ggplot2 (v. 3.4.2).

##### Pathway enrichment analysis

List of DEGs (FDR < 0.01, and logFC >= 1) was uploaded to Ingenuity Pathways Analysis (IPA) to get the list of enriched pathways. Raw data was downloaded from IPA into R and plotted using ggplot2 (v. 3.4.2). Only the top 20 pathways based on p-value were plotted. *Single Sample Gene Set Enrichment Analysis (ssGSEA)* ssGSEA was performed using normalized, log2 transformed expression data to calculate enrichment score (ES) using gsva() function from R Bioconductor package GSVA (v. 1.48.2) (Hänzelmann, Castelo, and Guinney 2013). Genes set to reflect enrichment of adherent junction was downloaded from Molecular Signatures Database (MSigDB). Genes set for epithelial mesenchymal transition was obtained from Liberzon et al (Liberzon et al. 2011).

### Statistical analysis

For statistical analysis and graphical presentation, Graphpad prism V10.1.0 (Domatics) software was used. Numerical results are given as means ± SD (N=sample size). The statistical significance for Western blot, CWB and Q-PCR was assessed with Graphpad with unpaired Student’s t test. For CD45+ cells present in the IN or OUT fraction, paired Student t-test was used for each cell line. The statistical significance for the comparison of genes expression and enrichment score was calculated using unpaired t test using R programming function “stat_compare_means” from ggpubr package. Statistical significance was accepted for *p<0.05; **p<0.01; ***p<0.001; ****p<0.0001.

## Results

### MUC2 differential expression between mucinous and non-mucinous hypermutated CRC is associated with low ICR score

In our analysis of the AC-ICAM cohort, we discerned a correlation between tumor histology (mucinous versus all other histology) and their immune infiltration and activation measured by their ICR score^1^. Differential gene expression analysis in mucinous adenocarcinoma vs. other adenocarcinoma showed that 3 mucin genes were over expressed in mucinous cancers. The most significant was MUC2 (p = 5.38e-16, logFC = 3.50), followed by MUC5B (p = 5.40e-11, logFC = 2.50), then MUC6 (p = 2.80e-9, logFC = 2.63) as illustrated in supplementary figure 4. This inverse correlation between the ICR score and the expression of MUC2 led us to perform functional analysis of the role of Mucin 2 in the immune infiltration of CRC in an *in-vitro* setting.

### MUC2 expression in CRC cell lines

In this context we analyzed the MUC2 expression in three commonly used CRC cell lines: LS-174T, HT-29 and Lovo by western blot and Q-PCR. As previously published^19^ in 2D culture LS-174T expressed MUC2 more significantly than HT-29, while expression in LoVo was undetectable (Figure 1A-C). A similar pattern of expression of MUC2 expression was observed in 3D culture of the same cell lines (Figure 1D-F). We therefore proceeded with the creation of MUC2 K.O. mutant in the 2 cell lines with MUC2 expression: HT-29 and LS-174T.

**Figure 1.**
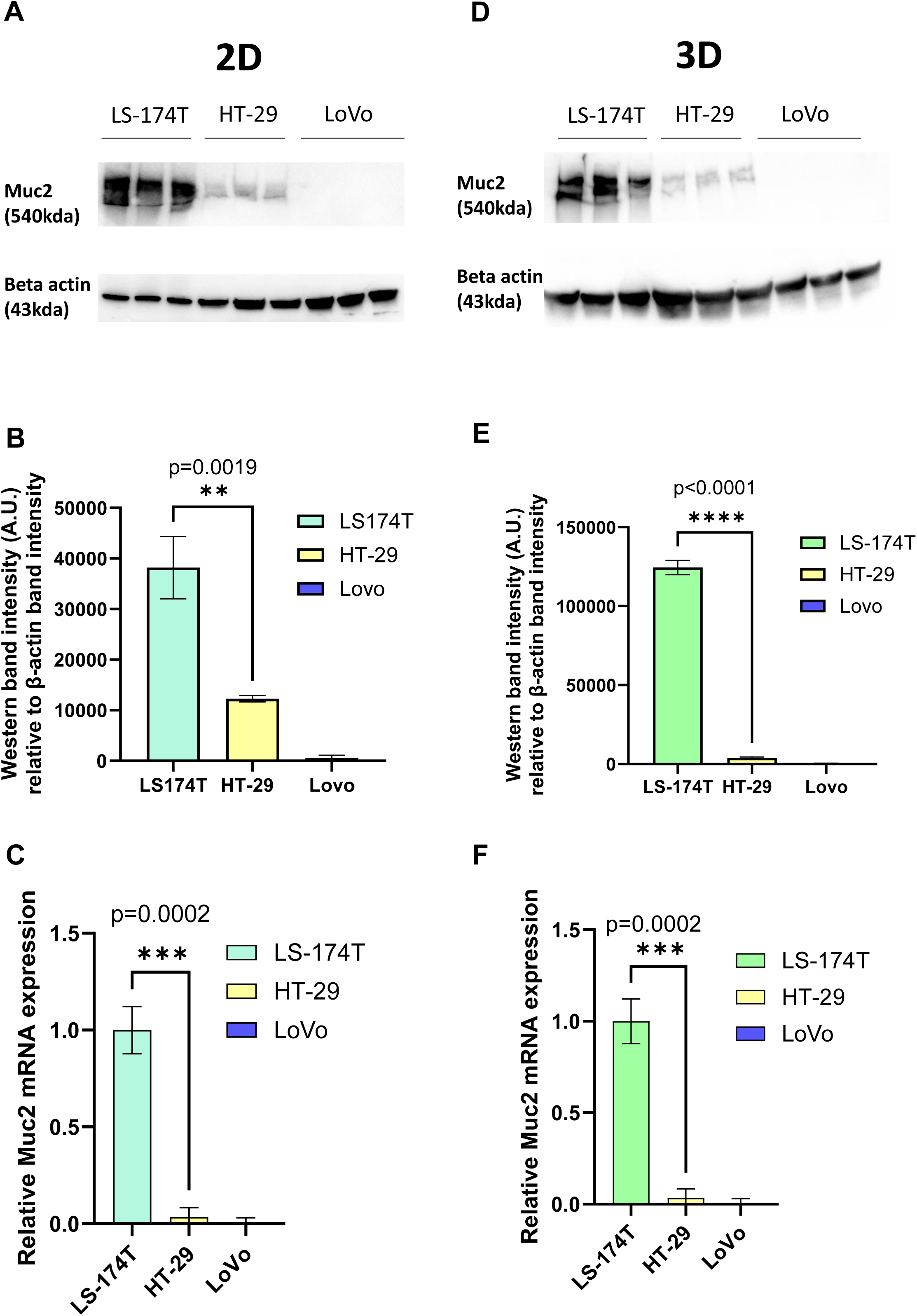
A. Westernblot of full length MUC2 protein (540kDa) and beta actin (housekeeping gene, 43kDa) in 3 different cell lines LS-174T, HT-29 and LoVo grown in 2D (N=3 biological replicates). B. Quantification of band intensity of MUC2 protein in those 3 cells lines grown in 2D. MUC2 is present in abundance in LS-174T, with significantly less expression in HT-29 (p=0.0019) (unpaired T-test) and no expression in LoVo cell line, is observed. C. Relative mRNA expression of MUC2 analyzed by Q-RT-PCR (N=3 biological replicates) in the 3 cell line grown in 2D. MUC2 expression is strongest in LS-174T compared to HT-29 (p=0.0002) (unpaired T-test) and no expression in LoVo is observed. D. Westernblot of full length MUC2 protein (540kDa) and beta actin (43 kDa) in 3 different cell lines LS-174T, HT-29 and LoVo grown in 3D (N=3 biological replicates). E. Quantification of band intensity of MUC2 protein in those 3 cells lines grown in 2D. MUC2 is present in abundance in LS-174T, with significantly less expression in HT-29 (p<0.0001) (unpaired T-test) and no expression in LoVo cell line, is observed. F. Relative mRNA expression of MUC2 analyzed by Q-RT-PCR (N=3 biological replicates) in the 3 cell line grown in 2D. MUC2 expression is strongest in LS-174T compared to HT-29 (p=0.0002) (unpaired T-test) and no expression in LoVo is observed.

### MUC2 Knock Out in HT-29 and LS-174T cell lines

5 gRNA were designed for K.O. of MUC2 targeting exon 2. Each gRNA was tested on each cell line by transfecting Cas9 ribonucleic complex by electroporation. TIDE analysis was performed for each gRNA on both cell lines. For HT-29 gRNA 2 was selected for further single cell cloning with an approximate efficiency of 55% (Supplementary Figure 5A). After single cell cloning and sanger sequencing of each clone and analysis with ICE, HT-29 gRNA2 clone 22 was selected for further work. For this clone the knockout was performed by introduction of 2 bases after the PAM sequence on both alleles inducing a frame shift and truncated protein production (supplementary figure 5B). This mutation was further validated by whole genome DNA sequencing (additional data).

For LS-174T, after TIDE analysis gRNA 3 was selected for single cell cloning with an approximate efficiency of 57% (Supplementary Figure 5C). After single cell cloning and sanger sequencing of each clone and analysis with ICE, LS-174T gRNA 3 clone 2 was selected for further work. For this clone the knockout was performed by introduction of 1 base after the PAM sequence on both alleles (Supplementary Figure 5D). This mutation was also further validated by whole genome DNA sequencing (additional data).

For both cell lines, to verify the specificity of the CRISPR edition and the absence of off target edition, whole genome sequencing (WGS) was performed and no potential off target mutations were detected in the 2 clones selected (supplementary data).

The knockout of MUC2 expression was further validated by western blot in 2D and 3D for both cell lines (Figure 2A/B). Additionally, the lack of MUC2 protein was validated by immune staining in 3D for both cell lines (figure 2C).

**Figure 2.**
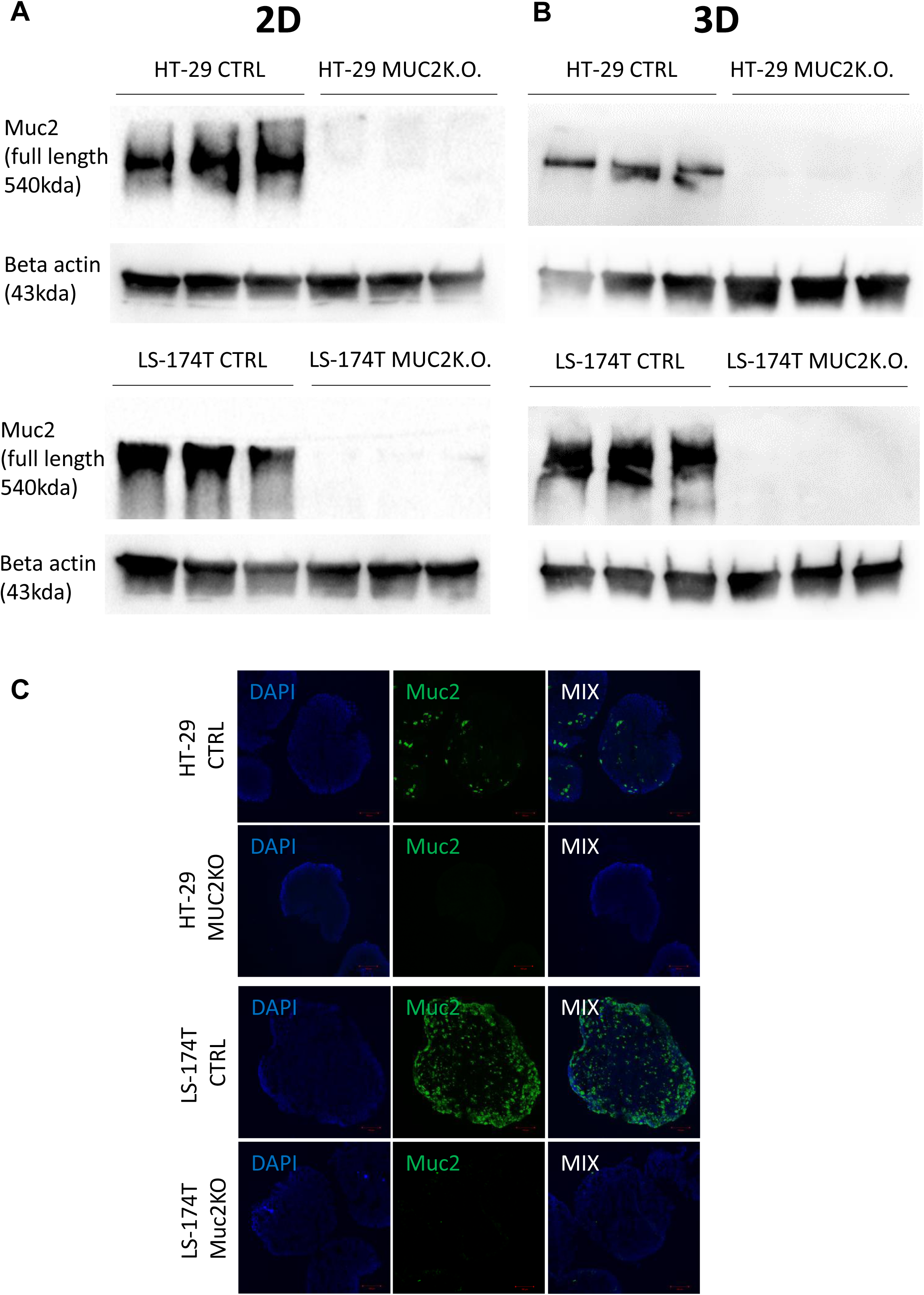
A. Westernblot of full length MUC2 protein (540kDa) and beta actin (43 kDa) in 3 different cell lines LS-174T CTRL, LS-174T MUC2 K.O., HT-29 CTRL and HT-29 MUC2 K.O. grown in 2D (N=3 biological replicates). B. Westernblot of full length MUC2 protein (540kDa) and beta actin (43 kDa) in 3 different cell lines LS-174T CTRL, LS-174T MUC2 K.O., HT-29 CTRL and HT-29 MUC2 K.O. grown in 2D (N=3 biological replicates). C. Representative images of spheroids of HT-29 and LS-174T CTRL and MUC2 K.O. immunostaining with Mouse anti-hMUC2 antibody (green) and counterstained with DAPI. Images acquired on confocal LS780. Scale bar of 100µm.

All those data together, sanger sequencing, WGS, western blot and immune staining demonstrated the knockout of the MUC2 protein in both cell lines HT-29 and LS-174T further designated as HT-29 MUC2 K.O. and LS-174T MUC2 K.O. To facilitate analysis and imaging those clones were further modified as well as their CTRL counterpart to express GFP as described in material and methods.

### Effect of MUC2 knock out on cellular behavior

We then further characterized our mutant cell lines in comparison to their control counterpart by measuring GFP using the GFP modified cell lines. In 2D culture, HT-29 MUC2 K.O. demonstrated a significantly slower growth than the original control counterpart (p=0.0011) (Figure 3A). In contrast, LS-174T MUC2 K.O. showed similar growth compared to the control counterpart (Figure 3B). On the other hand, when cultured in 3D HT-29 MUC2 K.O. showed no significant difference with the control cells (Figure 3C), however this time LS-174T MUC2 K.O. grew significantly slower than the control cells (p=0.0134) (Figure 3D). Additionally, we measured the size of the spheroids in 3D over 5 days of culture. Similarly, to the cell growth, HT-29 CTRL and HT-29 MUC2 K.O. displayed similar areas across 7 days (supplementary Figure 6A), while LS-174T CTRL spheroids area was higher than that of LS-174T MUC2 K.O. spheroids (Supplementary Figure 6B).

**Figure 3.**
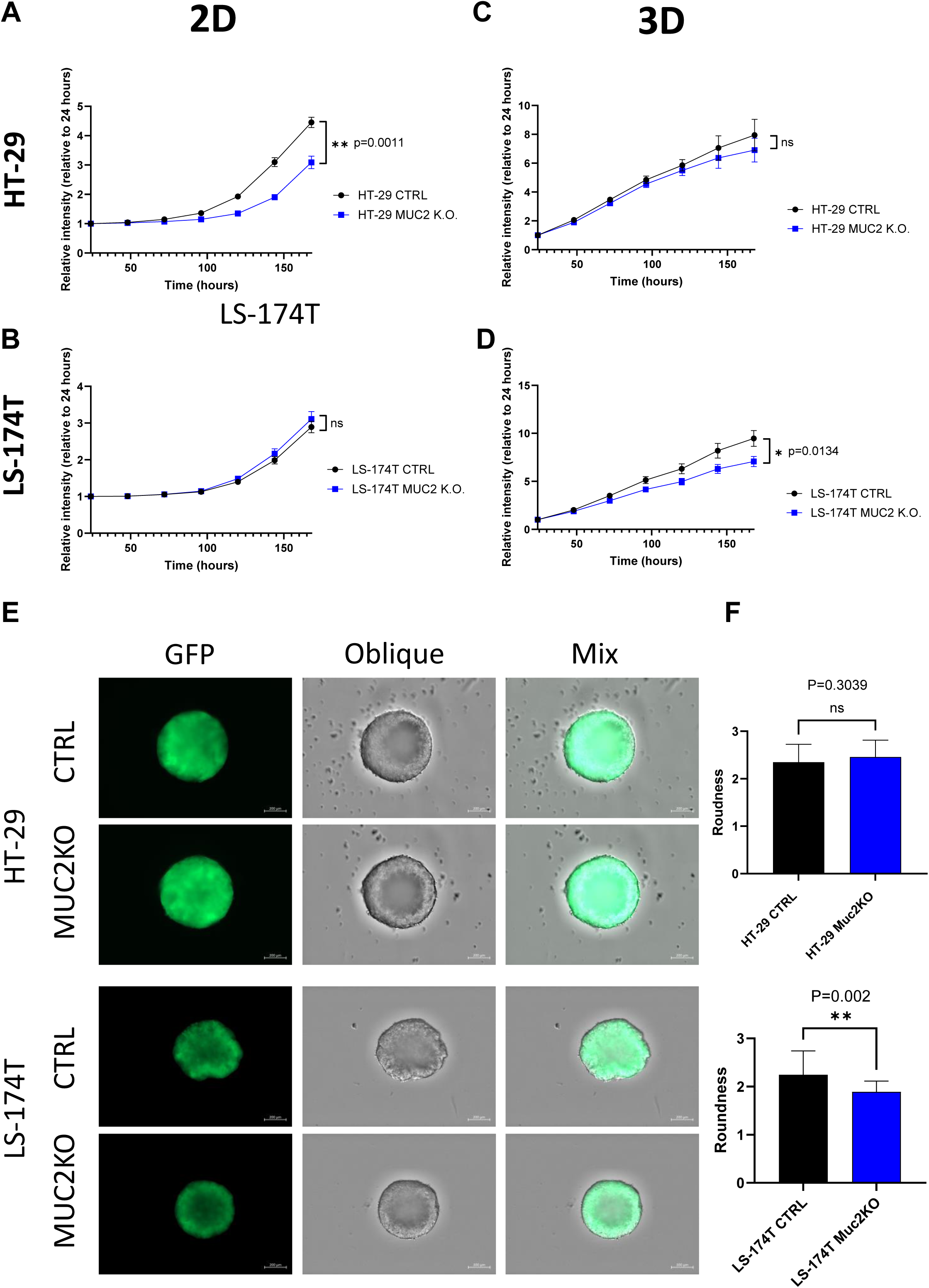
A. HT-29 CTRL and HT-29 MUC2 K.O. growth in 2D culture setup monitored by plate reader for 7 days by GFP intensity (N=3 biological replicates in 30 technical replicate each). HT-29 CTRL grow significantly faster than HT-29 MUC2 K.O. (p=0.0011) (unpaired t-test). B. LS-174T CTRL and LS-174T MUC2 K.O. growth in 2D culture setup monitored by plate reader for 7 days by GFP intensity (N=3 biological replicates in 30 technical replicate each). No significant difference was observed (unpaired t-test). C. HT-29 CTRL and HT-29 MUC2 K.O. growth in 2D culture setup monitored by plate reader for 7 days by GFP intensity (N=3 biological replicates in 30 technical replicate each). No significant difference was observed (unpaired t-test). D. LS-174T CTRL and LS-174T MUC2 K.O. growth in 2D culture setup monitored by plate reader for 7 days by GFP intensity (N=3 biological replicates in 30 technical replicates each). LS-174T CTRL grew significantly faster than LS-174T MUC2 K.O. (p=0.0134) (unpaired t-test). E. Representative images of day 5 spheroids acquired with zeiss CD7 fluorescent microscope of GFP and oblique of spheroids of HT-29 and LS-174T CTRL or MUC2 K.O. Scale of 200µm. F. Roundness analysis of HT-29 (N=13 each, biological replicates) and LS-174T (N=25 each, biological replicates) of CTRL and MUC2 K.O. performed by analysis with Zen blue software of area and perimeter of each spheroid. No significant difference is observed between HT-29 CTRL and HT-29 MUC2 K.O., but LS-174T MUC2 K.O. is significantly more round than LS-174T CTRL (unpaired t-test).

Incidentally, LS-174T CTRL could form spheroids with secondary appendix in 3D culture while HT-29 CTRL spheroids appeared to be rounder and smoother (Figure 3E). We can note that LoVo cells could barely form spheroids after 5 days of culture (data not shown). Upon knock out of MUC2, the 3D morphology was not significantly affected in HT-29 (Figure 3F), while the roundness was significantly increased in LS-174T MUC2 K.O. compared to CTRL (p=0.002) (Figure E-F) and with secondary appendix far less noticeable. The single sample gene set enrichment analysis (ssGSEA) of HT-29 CTRL against HT-29 MUC2 K.O. as well as LS-174T CTRL against LS-174T MUC2 K.O. showed that while we could see an increase in the adherence junction enrichment score for the HT-29 MUC2 K.O. (p = 0.041), we observed a very strong decrease of the enrichment score in LS-174T upon K.O. of MUC2 (p = 0.047) (Supplementary Figure 7A). This opposite effect observed on the adherence junction genes could explain in part the opposite change in morphology of the two cell lines and potentially also the changes in cell proliferation in the 2D and 3D models.

As it was previously demonstrated that MUC2 silencing promoted CRC metastasis^19^ we investigated whether we could observe increased migration or upregulated EMT markers in either of our cell lines. While we could observe cell proliferation, little to no migration was observed in HT-29 cancer cell lines with or without MUC2 even after 72h. In both cases no increase in migration was observed in MUC2 K.O. compared to CTRL when tested by scratch test (Supplementary Figure 8 A-D) and limited if any migration was observed by transwell (data not shown). We also investigated the EMT markers such as E-Cadherin, N-cadherin and Vimentin by western blot in our control and MUC2 K.O. (Supplementary Figure 7 B-E) and observed no difference in their protein expression with or without presence of MUC2 protein expression. We also investigated EMT genes signature by RNA expression between CTRL and MUC2 K.O. in both cell lines and could not identify any increase in HT-29 and even saw a minimal but significant decrease in LS-174T upon knockout of MUC2 both in 2D (p = 0.00041) and 3D (n = 0.046)(Supplementary Figure 9).

In summary, the effects of MUC2 knockout differed between the two cell lines. HT-29 cells with MUC2 knockout grew slower in 2D culture, indicating a possible regulatory function of MUC2 in their proliferation. On the other hand, LS-174T cells with MUC2 knockout showed similar growth in 2D but slower growth in 3D culture, implying that the impact of MUC2 on cell behavior may depend on the environment.

### Effect of MUC2 knockout on killing by allogenic enriched activated PBMCs

We first investigated if the absence of MUC2 protein would give better access to being killed by allogenic activated T cells from PBMCs enriched in T cells and NK cells (E/A PBMCs). In a 2D setup we plated 500 GFP expressing cancer cells and allowed them to adhere for 48h prior to the addition of varying numbers of A/E PBMCs. The number of cancer cells was monitored by plate reader for up to 5 days of co-culture by GFP measurement. To account for the difference in growth speed documented previously between CTRL and MUC2 K.O., the results were normalized to control cells without PBMCs. Similarly in a 3D setup we plated 2000 GFP expressing cancer cells in ultralow adherence 96 well plates and allowed them to form spheroids for 5 days prior to the addition of varying numbers of A/E PBMCs. The number of cancer cells was monitored by GFP expression measured on a plate reader for up to 4 days of co-culture. While both in 2D and 3D we could observe a decrease in cancer cells with every increase in the amount of E/A PBMCs plated on the cancer cells, no significant difference could be observed between CTRL and MUC2 K.O. in both cell lines (supplementary figure 10 A - D). Though, we could identify a higher sensitivity of HT-29 cells compared to LS-174T in both CTRL and MUC2 K.O. to co-culture with 500 A/E PBMCs in 2D and co-culture with 500 A/E PBMCs in 3D (Figure 4A). Representative video of the co-culture in 3D over 2 days are shown as supplementary data (Supplementary Figure 11). It shows a strong interaction of the PBMCs and a strong expansion of the E/A PBMCs when co-cultured with cancer cells. Yet no drastic visual differences between CTRL and MUC2 K.O. could be observed. Using the plate readers measurement of the whole wells with GFP cancer cell could not demonstrate an increase in cancer cell killing by removal of MUC2 protein in either of the cell lines, using any of the E/A PBMC numbers, in either a 2D or 3D setting. Overall, using basic methods such as plate readers, we were unable to discern any heightened efficacy in the killing of MUC2 knockout cells by enriched activated PBMCs, neither in 2D nor in 3D settings. Only in microscopy after rinse on the spheroids we could see an effect on HT-29 MUC2 K.O.

**Figure 4.**
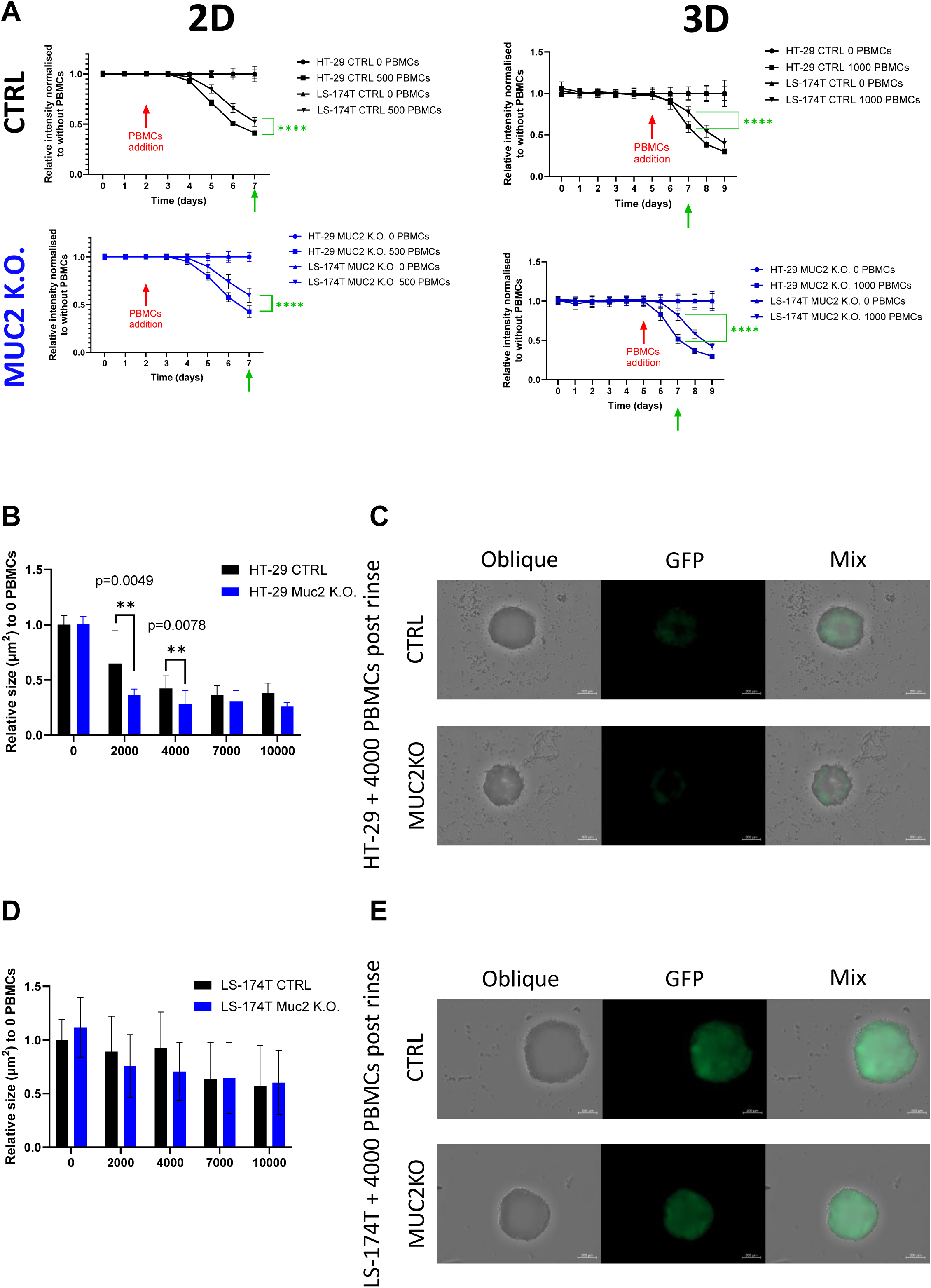
A. HT-29 and LS-174T, CTRL and MUC2 K.O. growth and death monitored by GFP expression over 7 days in 2D and 9 days in 3D culture set up. 0 or 500 E/A PBMCs were added on day 2 for 2D and 0 or 1000 E/A PBMCs were added on day 5 for 3D co-culture setup. Data are presented as relative to the same cell line without co-culture with PBMCs (0 E/A PBMCs) (N=18, 3 biological replicates of 6 technical replicates). Significant differences between HT-29 and LS-174T cell lines were calculated at day 7 in 2D and 3D by student T-test. B. HT-29 CTRL and MUC2 K.O. spheroids area of IN fraction spheroids post rinse after co-culture for 2 days with E/A PBCMs relative to area of IN fraction after rinse without co-culture. After co-culture with 2000 or 4000 PBMCs area of IN fraction spheroids is significantly smaller in HT-29 MUC2 K.O. than in HT-29 CTRL (N=12, 2 biological of 6 technical replicates) (unpaired t-test). C. Representative images of IN fraction spheroids post rinse after co-culture with 4000 E/A PBMCs of HT-29 CTRL and HT-29 MUC2 K.O. Images were acquired for oblique and GFP expression on Zeiss CD7, and area analyzed with Imaris software. Scale bar of 200µm. D. LS-174T CTRL and MUC2 K.O. spheroids area of IN fraction spheroids post rinse after co-culture for 2 days with E/A PBCMs relative to area of IN fraction after rinse without co-culture. No significant difference was observed between LS-174T CTRL and MUC2 K.O. at any number of E/A PBMCs co-culture (N=12, 2 biological of 6 technical replicates) (unpaired t-test). E. Representative images of IN fraction spheroids post rinse after co-culture with 4000 E/A PBMCs of LS-174T CTRL and LS-174T MUC2 K.O. Images were acquired for oblique and GFP expression on Zeiss CD7, and area analyzed with Imaris software. Scale bar of 200µm.

### Effect of MUC2 Knock Out on PBMCs infiltration

We investigated by microscopy and flow cytometry the E/A PBMCs infiltration of the spheroids. Using the same conditions previously discussed we co-cultured E/A PBMCs with cancer cells. The IN and OUT fraction were then separated and the remaining IN fraction size was evaluated by microscopy. A significant size difference was observed with HT-29, with smaller IN fraction left with HT-29 MUC2 K.O. compared to CTRL after 2 days of co-culture with 2000 PBMCs (p=0.0049) or 4000 PBMCs (p=0.0078) (Figure 4B-D), while no significant difference was observed between the remaining IN fraction of LS- 174T CTRL and LS-174T MUC2 K.O. (Figure 4D-E).

To further investigate the number of cells within each fraction we performed flow cytometry analysis of the 2 fractions for each cell line. When analyzing by flow cytometry the IN fraction in absence of co-culture with PBMCs, we noticed that we could recoup significantly less Epcam+ cells in HT-29 MUC2 K.O. than in HT-29 CTRL (p=0.0182) (Figure 5A) despite documenting previously similar growth in 3D setup between CTRL and MUC2 K.O (Figure 3C and Supplementary Figure 6A). The increase in this cell line of adherens junction expression (Supplementary Figure 7A) could explain in part this difference. Also, we noticed that during filtration on 40µm cell strainer prior to flow cytometry analysis more clumps were retained with HT-29 MUC2 K.O spheroids than their CTRL counterpart (data not shown). On the other hand, with LS-174T, even if not significant we tend to recover slightly more Epcam+ cells in LS-174T MUC2 K.O. in fraction than in the CTRL counterpart. Once again this might be due to the reduction in expression of adherens junction proteins between the cells in the MUC2 K.O. compared to the CTRL in this cell line, facilitating more efficient dissociation. A direct comparison of the number of Epcam+ cells recovered in the IN fraction was therefore not possible. The CD45+ cells on the other hand should not have been affected by this adherens junction expression in the cancer cells and we could compare the number of CD45+ cells recovered. We saw a significant increase of CD45+ cells in the IN fraction of HT-29 MUC2 K.O. compared to CTRL (Figure 5B). In contrast, no difference was observed in LS-174T. Also, in the OUT fraction, even if not significant we tended to recover more CD45+ cells after co-culture with HT-29 MUC2 K.O. than with the CTRL cells. Inversely slightly less CD45+ cells were recovered with LS-174 MUC2 K.O. than with CTRL.

**Figure 5.**
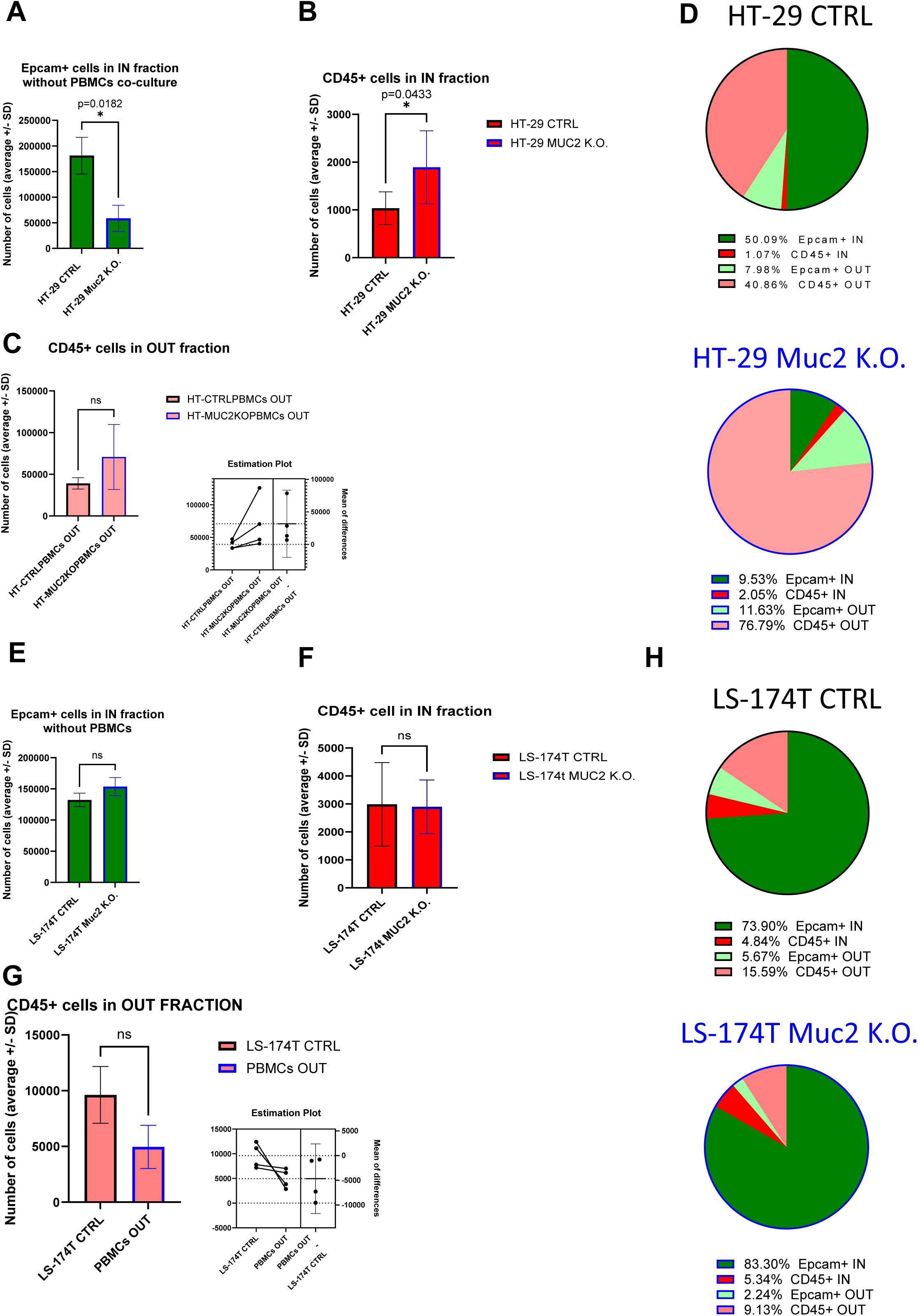
A. Average number of Epcam+ cells recovered in the IN fraction after digestion, staining and flow analysis without co-culture with E/A PBMCs. Significantly less Epcam+ cells were recovered for HT-29 CTRL than for HT-29 MUC2 K.O. (p=0.0182, Paired T-test) (N=4 biological replicates). B. Average number of CD45+ cells recovered in the IN fraction after digestion, staining and flow analysis. Significantly less CD45+ cells were recovered for HT-29 CTRL than for HT-29 MUC2 K.O. (p=0.0433, Paired T-test) (N=4 biological replicates). C. Average number of CD45+ cells recovered in the OUT fraction after digestion, staining and flow analysis. No significant difference in CD45+ cells recovered after co-culture with HT-29 CTRL or HT-29 MUC2 K.O. was detected despite having systematically more CD45+ cells after co-culture with HT-29 MUC2 K.O. than with HT-29 CTRL (p=0.1460, Paired T-test) (N=4 biological replicates). D. Pie chart representation of average relative abundance of Epcam+ and CD45+ cells in the IN and OUT fraction after 2 days of co-culture for HT-29 CTRL and MUC2 K.O. (N=4 biological replicates). E. Average number of Epcam+ cells recovered in the IN fraction after digestion, staining and flow analysis without co-culture with E/A PBMCs. No statistical difference was observed in Epcam+ cells recovered between LS-174T CTRL and LS-147T MUC2 K.O. (p=0.1405, Paired T-test) (N=3 biological replicates). F. Average number of CD45+ cells recovered in the IN fraction after digestion, staining and flow analysis. No difference in CD45+ cells recovered was observed between LS- 174T CTRL and LS-174T MUC2 K.O. (p=0.9202, Paired T-test) (N=4 biological replicates). G. Average number of CD45+ cells recovered in the OUT fraction after digestion, staining and flow analysis. No significant difference in CD45+ cells recovered after co-culture with LS-174T CTRL or LS-174T MUC2 K.O. was detected despite having systematically more CD45+ cells after co-culture with LS-174T CTRL. than with LS-174T MUC2 K.O. (p=0.1262, Paired T-test) (N=4 biological replicates). H. Pie chart representation of average relative abundance of Epcam+ and CD45+ cells in the IN and OUT fraction after 2 days of co-culture for LS174T CTRL and MUC2 K.O. (N=4 biological replicates).

In summary, our flow cytometry analysis revealed that the knockout (K.O.) of MUC2 in HT-29 cells resulted in heightened proliferation of E/A PBMCs and an augmentation in immune cell infiltration in the cancer cell spheroids, resulting in more cancer cell elimination and reduction in cancer spheroids. Conversely, such effects were not evident in LS-174T cells. Together This comes as a confirmation of the observation previously made in microscopy (Figure 4 B-E)

### DEG analysis of the invading PBMCs show cell cycle and Interferon pathway activation in HT-29 MUC2 K.O. cells only

To further understand the different impact MUC2 knock out has on our two cell lines we performed RNA sequencing analysis of both cell lines with and without co-culture. The RNA seq was performed in 2D and 3D for CTRL and MUC2 K.O. cells, in addition to CTRL PBMCs (each RNA seq was performed in biological triplicates). Global transcriptional change was assessed using PCA. Results show a clear separation between CTRL-3D (green) and MUC2 K.O.-3D (brown)) in both cell lines. In HT-29 cell line, CTRL + E/A PBMCs IN fraction (blue) separates from MUC2 K.O. + E/A PBMCs IN fraction (purple), while this much less so for LS-174T. Moreover, there is a separate cluster of CTRL PBMCs (peach), CTRL E/A PBMCs D2 (control enriched/activated PBMCs at day 2) (brown) and CTRL E/A PBMCs D7 (red) (Figure 6A).

**Figure 6.**
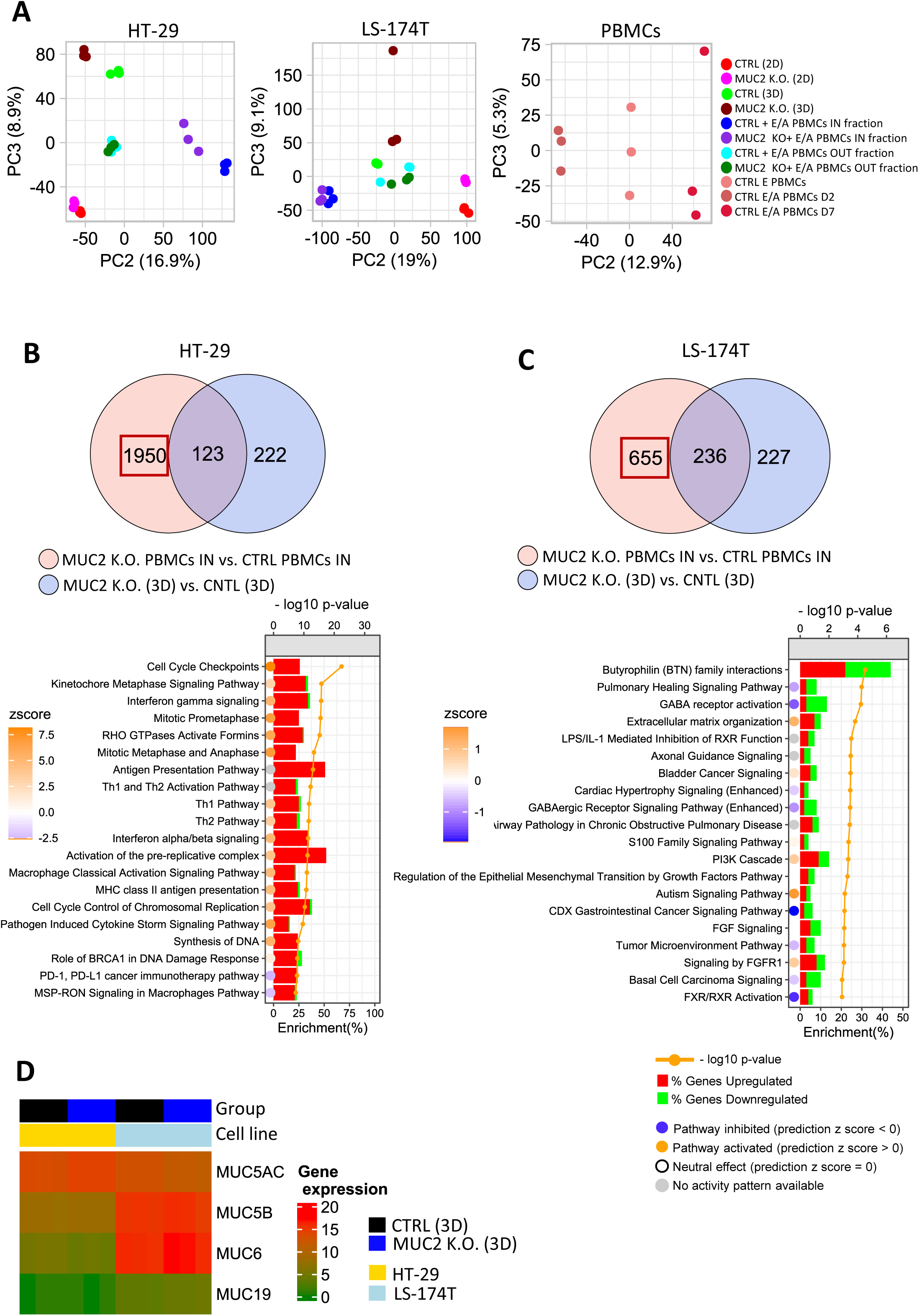
A. Principal component analysis (PCA) based on gene expression profile in HT-29, LS- 174T cultured in 2D and 3D, E/A PBMCs or IN and OUT fraction of co-culture of cancer cells with E/A PBMCs. B: Venn diagram of DEG (FDR < 0.01, logFC >= 1) between HT-29 MUC2 K.O. + E/A PBMCs IN Fraction vs. HT-29 CTRL + E/A PBMCs IN fraction (n = 2073 genes) and HT- 29 MUC2 K.O. (3D) vs. HT-29 CTRL (3D) (n = 345). Pathways enrichment analysis was performed on genes reflecting DEG between PBMCs IN (MUC2) vs. PBMCs IN (CTRL) (n = 1950) by subtracting the common genes. Top 20 pathways based on p-value were selected for plotting. Comparison of those two DEG lists allows us to see that when in contact with HT-29 MUC2 K.O., E/A PBMCs increase their cell cycle and are IFN pathway activation. C. Venn diagram of DEG (FDR < 0.01, logFC >= 1) between LS-174T MUC2 + E/A PBMCs IN fraction vs. LS-174T CTRL + E/A PBMCs IN fraction (n = 891) and MUC2 K.O. (3D) vs. CNTL (3D) (n = 463). Pathways enrichment analysis was performed on genes reflecting DEG between PBMCs IN (MUC2) vs. PBMCs IN (CTRL) (n = 655) by subtracting common the common genes. Top 20 pathways based on p-value were selected for plotting (FDR < 0.01). Comparison of those two DEG lists does not demonstrate similar pattern with LS-174T. D. Genes expression of other gel forming mucin in HT-29 and LS-174T spheroids shows that LS-174T also have strong expression of MUC5B, MUC6 and MUC19 compared to HT-29. A and B. Histograms represent the proportion (%) of DEGs upregulated (red) or downregulated (green) in PBMCs IN (MUC2) vs. PBMCs IN (CTRL). The circles represent the pathway activation status. Blue circle indicates the pathway is inhibited with a negative z-score, orange circle represents a pathway is activated with a positive z-score, the white circle represents the pathway is neutral with zero z-score, while a gray circle indicates that the pathway activity is unknown.

In conclusion, PCA analysis revealed distinct global transcriptional changes due to MUC2 knockout in both HT-29 and LS-174T cell lines, with PCA showing clear separation between 3D cultures of CTRL and MUC2 K.O. cells. More importantly, HT-29 cells co-cultured with PBMCs showed significant separation between CTRL and MUC2 K.O. conditions, unlike LS-174T, highlighting cell line-specific responses to PBMCs co-culture when MUC2 is knocked out.

We first evaluated the effect MUC2 KO had on each cell line.HT-29 cells grown in 2D displayed upon MUC KO, significant upregulation of genes in pathways associated with cell cycle checkpoints. This is compatible with our observation of significantly slower cell proliferation in cells missing MUC2 expression. (Supplementary Figure 13A). In 3D little difference in gene expression was observed for HT-29 (Supplementary Figure 13B). In LS-174T cells, while a significant number of DEGs were identified in both in 2D and 3D, no particular or significant p-value enriched pathways were noticed (Supplementary Figure 13 C-D).

In order to then compare the co-cultures we first performed the RNA sequencing of the PBMCs alone before and after activation prior to co-culture with cancer cells. We then performed RNA sequencing of the IN (part of the spheroid) and OUT (in suspension) fraction of each cell line after 2 days of co-culture. To identify genes specific to PBMCs in the cell fractions, we compared DEGs between CTRL cancer cells + PBMCs and MUC2 K.O. cancer cells + PBMCs, then intersected this list with DEGs from comparing CTRL cancer cells alone to MUC2 K.O. cells alone in 3D culture. By removing overlapping genes, we isolated genes likely differentially expressed by PBMCs after co-culture or by cancer cells after PBMC contact (Figure 6B). In the HT-29 cells KO of MUC 2 followed by co-culture with PBMCs the identified DEGs (n = 1950) are involved with cell cycle checkpoints (-log10 p-value = 22.4, (p = 3.98e-23)), Kinetochore Metaphase Signaling Pathway (cycle progression pathway) (-log10 p-value = 15.8, (p = 1.58e-16)) and INF-γ signaling (-log10 p-value = 15.7, (p = 1.99e-16)) (Figure 6B). When the same analysis was performed with LS-174T cancer cells (n = 655) no significant pathway with strong p-value could be detected (Figure 6C). Also, when analyzing the ICR score in the IN co-culture fraction, only in HT-29 we noticed a significant increase of the ICR score (p=0.009) (Supplementary Figure 14)

While the presence of MUC2 mRNA is seen, due to the K.O. by indel it is not translated into functional protein in the K.O. cells (no protein detected by westernblot as verified previously), we verified the expression of other mucin genes known to be gel forming: MUC5AC, MUC5B, MUC6 and MUC19. MUC5AC expression was increased in HT-29 MUC2 K.O. compared to CTRL and decreased in LS-174T MUC2 K.O. MUC5B, was not significantly differentially expressed in both cell lines. However, MUC6 was significantly upregulated in CTRL compared to MUC2 K.O. in HT-29 (p = 0.023), whereasMUC19 was significantly differentially expressed between CTRL and MUC2 K.O. in LS-174T (p = 0.0036). When comparing the gene expression of MUC5B, MUC6 and MUC19 between the two cell lines, they were significantly more expressed in LS-174T cells than in HT-29 cells (Figure 6D and Supplementary Figure 15).

In summary, in MUC2 knockout cells, we saw and overall higher expression of MUC5B, MUC6, and MUC19 in LS-174T compared to HT-29.

## Discussion

The research conducted in this study aimed to unravel the intricate role of MUC2, a mucin protein, in the context of colorectal cancer (CRC), with a specific focus on its influence on immune response dynamics and cellular behavior. The investigation stemmed from the observation of an inverse correlation between MUC2 expression and the immune constant of rejection (ICR) score in mucinous adenocarcinomas compared to other histological types.

In interpreting our findings from the previously published multi-omics dataset comparing mucinous and non-mucinous CRC with the ICR, two hypotheses emerge. The first suggests that mucin may serve as a barrier against bacterial invasion, akin to its role in the normal colon mucosa. Subsequently deterring immune cell attraction by the intratumorally microbiome. However, our findings in the AC-ICAM study showed that there are bacterial genera associated with ICR and there are bacteria associated prognosis, these genera were not the same.

The second hypothesis, which is not mutually exclusive with the first, posits that mucin could act as a physical barrier directly impeding immune cells from penetrating the tissues. This study focuses on this second hypothesis and seems to validate these observations in vitro. Specifically, the removal of MUC2 in HT-29 cells resulted in heightened immune infiltration and enhanced allogenic recognition of cancer cells.

The study employed a multi-pronged approach, utilizing MUC2 knockout (K.O.) in CRC cell lines HT-29 and LS-174T, shedding light on the nuanced effects of MUC2 on various cellular processes. The successful knockout of MUC2 was meticulously validated through a series of analyses, including sanger sequencing, whole-genome DNA sequencing, Western blot, and immune staining. This rigorous validation process ensured the reliability of subsequent findings and conclusions. One of the intriguing observations was the divergent impact of MUC2 knockout on the two distinct cell lines. HT-29 MUC2 K.O. cells exhibited slower growth in 2D culture, hinting at a potential regulatory role of MUC2 in the proliferation of these cells or their interaction with the vessel. In contrast, LS-174T MUC2 K.O. cells demonstrated comparable growth in 2D but significantly slower growth in 3D culture, suggesting a context-dependent influence of MUC2 on cellular behavior. Further exploration into the molecular mechanisms revealed an opposite effect on tight junction genes in the two cell lines. This differential gene expression may underlie the observed variations in cell morphology and growth patterns, emphasizing the complexity of mucin involvement in CRC.

Importantly, the study delves into the impact of MUC2 on the immune response. Sensitive techniques such as quantitative microscopy and flow cytometry analyses unveiled a significant increase in immune cell infiltration in HT-29 MUC2 K.O. We also observed an increased activation and proliferation of the T cells as they seem to have better access to the HT-29 cells leading to increase allogeneic immune rejection. This strongly suggests that MUC2 acts as a physical barrier hindering immune cell recognition and activation.

The RNA sequencing data provided further insights, indicating that pathways associated with cell cycle and IFN-y signaling were upregulated in PBMCs co-cultured with HT-29 MUC2 K.O. Our data also shows an upregulation of the cell cycle checkpoints. We believe that this signature is due to the cancer cells in this remaining IN fraction as it was previously demonstrated that IFNs can indeed affect cell proliferation in tumor cells both by prolonging or blocking the cell cycle ^20–23^. This finding aligns with the enhanced immune infiltration observed in these conditions, reinforcing the notion that MUC2 impedes immune cell access to cancer spheroids.

The study also considered the expression of other gel-forming mucins, revealing that while Muc5A expression increased in HT-29 MUC2 K.O., Muc5B, Muc6, and Muc19 showed no significant differences and were significantly less expressed in HT-29 than in LS-174T. Therefore, we hypothesis that the absence of increased immune infiltration observed in LS-174T MUC2 K.O. is due to the presence of those other Mucin proteins in the ECM, in line with the presence of Muc5B and Muc6 significantly over expressed (with lower fold change and p-value than MUC2) genes in mucinous adenocarcinoma compared to other carcinomas.

In conclusion, this research contributes valuable insights into the multifaceted role of MUC2 in CRC. The divergent effects observed in HT-29 and LS-174T cells underscore the importance of considering cellular context and heterogeneity in studying mucin functions. These observations suggest that MUC2, and potentially other mucins, play a role in creating a barrier that limits immune cell access to cancerous tissues. This barrier function may prevent immune cell recognition and activation within the tumor microenvironment. The results of our study highlight the complex interplay between mucins, immune response, and the intricate dynamics of the tumor microenvironment in colorectal cancer. Further exploration of these mechanisms is warranted to elucidate the specific molecular pathways involved and to inform potential therapeutic strategies for manipulating mucin-related interactions in cancer immunity.

---

**Supplementary Figure 1.**
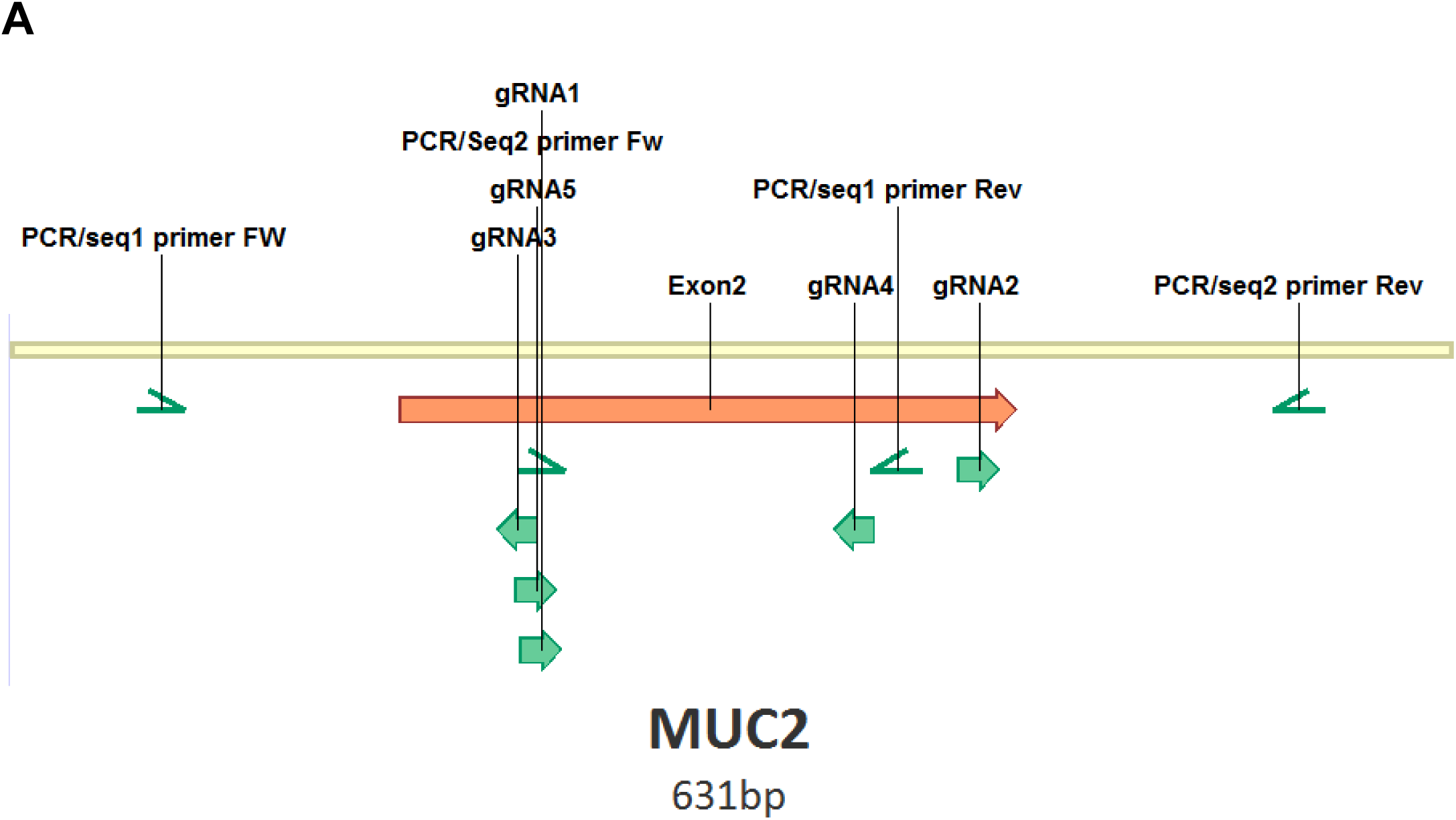
Relative localization and orientation of the 5 gRNAs design for Knock Out of MUC2 on Exon 2 and the primers used for PCR and sanger sequencing.

**Supplementary Figure 2.**
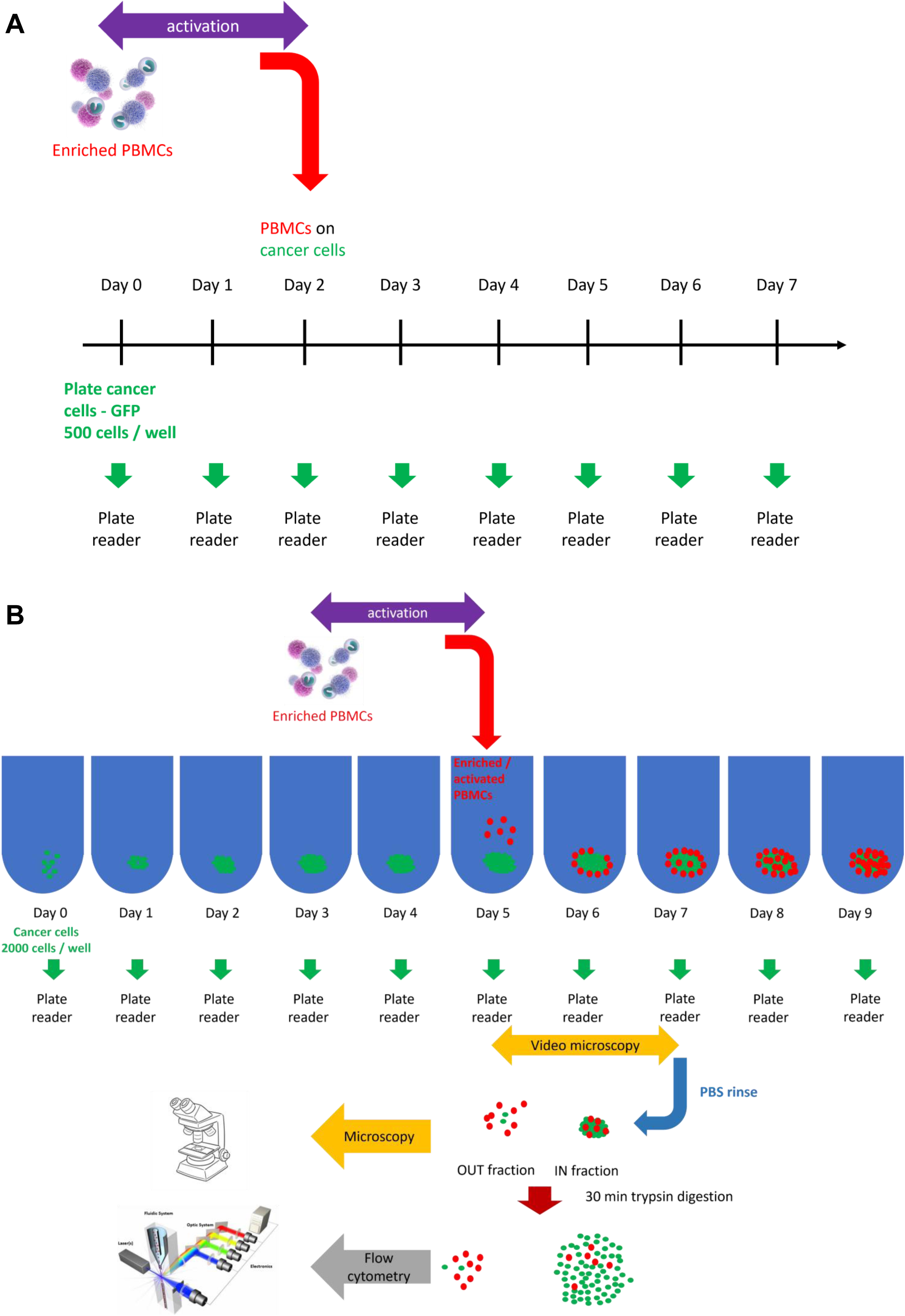
A. Schematic of the co-culture and analysis procedure used for 2D experiments. 500 cancer cells were seeded on each well of 96 well plate at day 0, allowed to settle and grow for 2 days. On day 0 enriched PBMCs were thawed and activated for 2 days. Various numbers of Enriched / activated cell are then seeded on top of the cancer cells at day 2. GFP expression is recorded every day from day 0 to day 7 on Ensight plate reader. B. Schematic of the co-culture and analysis procedure used for 3D experiments. 2000 cancer cells per well are seeded on low adherence U bottom well plates. Spheroids are allowed to be formed for 5 days. On day 3 enriched PBMCs were thawed and activated for 2 days. Co-culture was then performed for 2-4 days. GFP expression was monitored on plate reader from day 0 to day 9. Video-microscopy of co-culture was performed between day 5 and day 7. If microscopy analysis of remaining IN fraction is performed, spheroids were collected at day 7. For flow cytometry IN and OUT fractions are separated, then for flow cytometry each fraction is treated with trypsin prior staining and analysis.

**Supplementary Figure 3.**
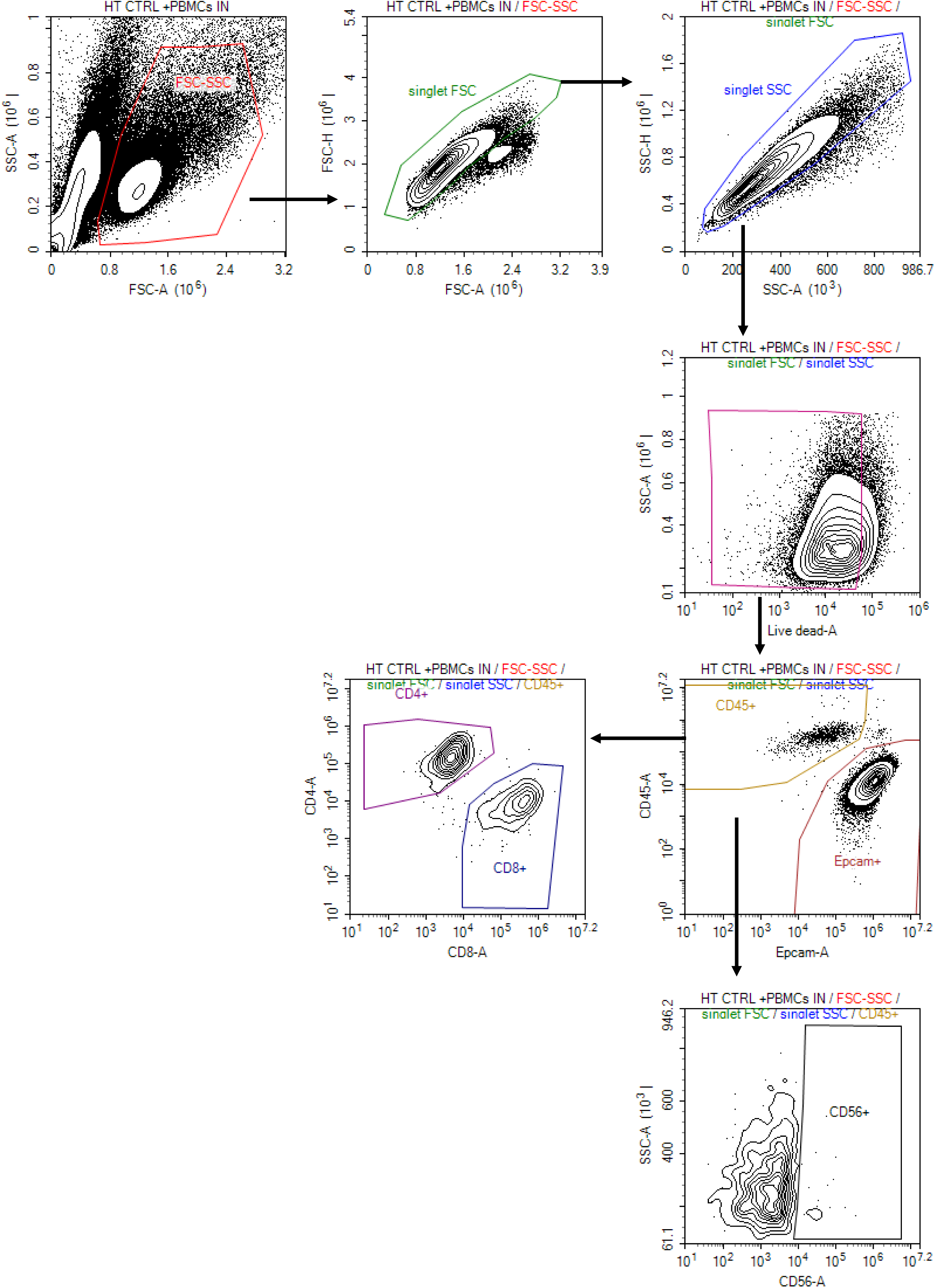
Gating strategy for flow cytometry analysis. Cells were first selected on FSC-A/SSC-A parameters. Singlets were selected on SSC-H/SSC-A and FSC-H/FSC-A parameters. Live cells were selected on the base of expression of SSC-A/ V2-A (Ex 405-Em530/30). Epcam+ and CD45+ cells were then selected from living cells based on B1-A / B2-A (Ex 488-Em615/24). CD45+ cells were further distinguished for CD4+/CD8a+ expression based on V1-A (Ex405-Em445/45) / V4-A (Ex405-Em615/24) and CD56+ cell were analyzed based on SSC-A / B4-A (Ex488-Em615/24).

**Supplementary Figure 4.**
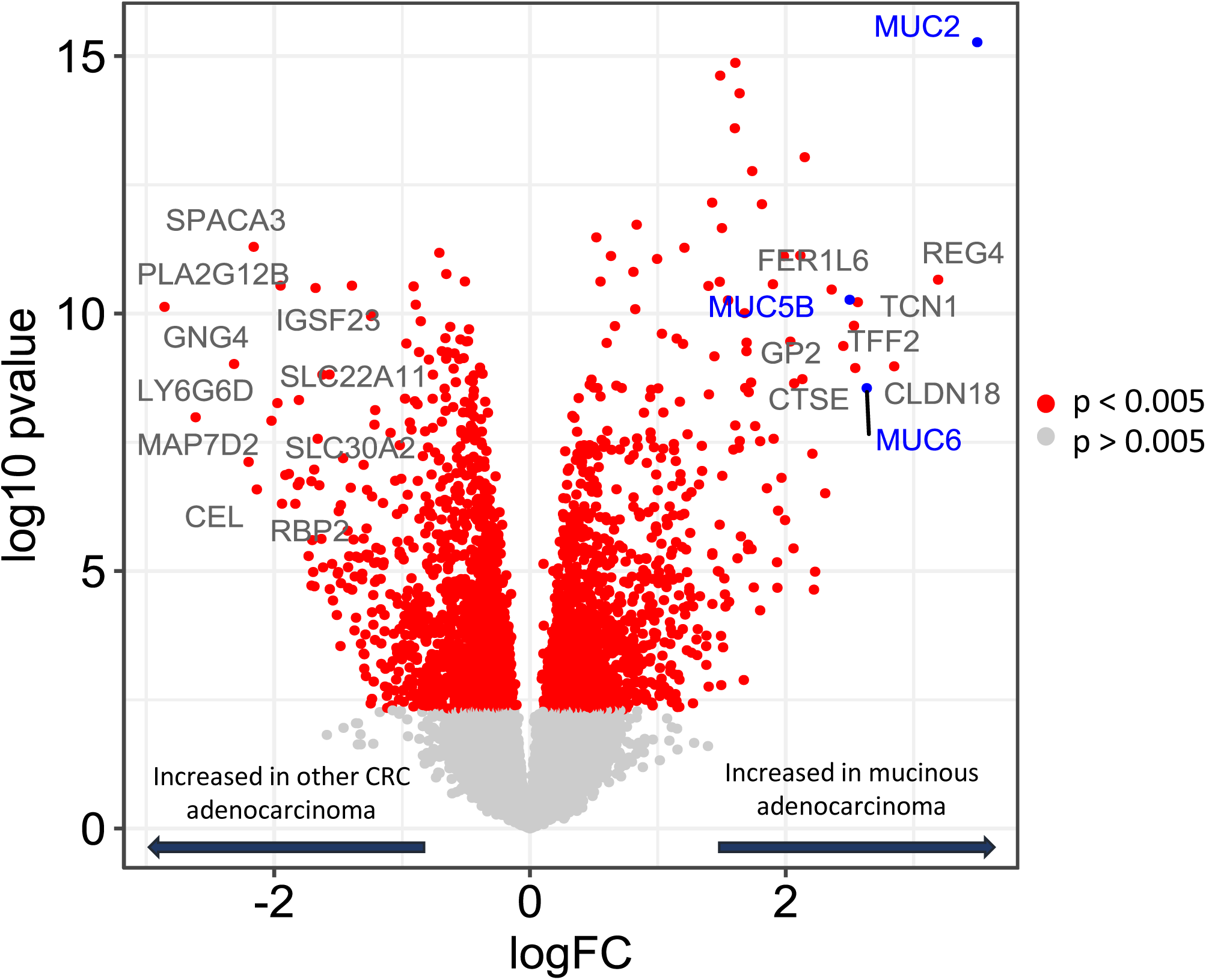
Volcano plot of the DEG using limma between mucinous versus all other histological classifications in hypermutated samples of the AC-ICAM cohort. The most significant gene expressed in mucinous samples with the higher fold change is MUC2 (p = 5.38e-16, logFC = 3.50). Two other mucins are identified as upregulated in mucinous samples: MUC5B (p = 5.40e-11, logFC = 2.50) and MUC6 (p = 2.80e-9, logFC = 2.63). Red dotes represents genes with p-value < 0.005 and grey dotes for genes with p-value > 0.005.

**Supplementary Figure 5.**
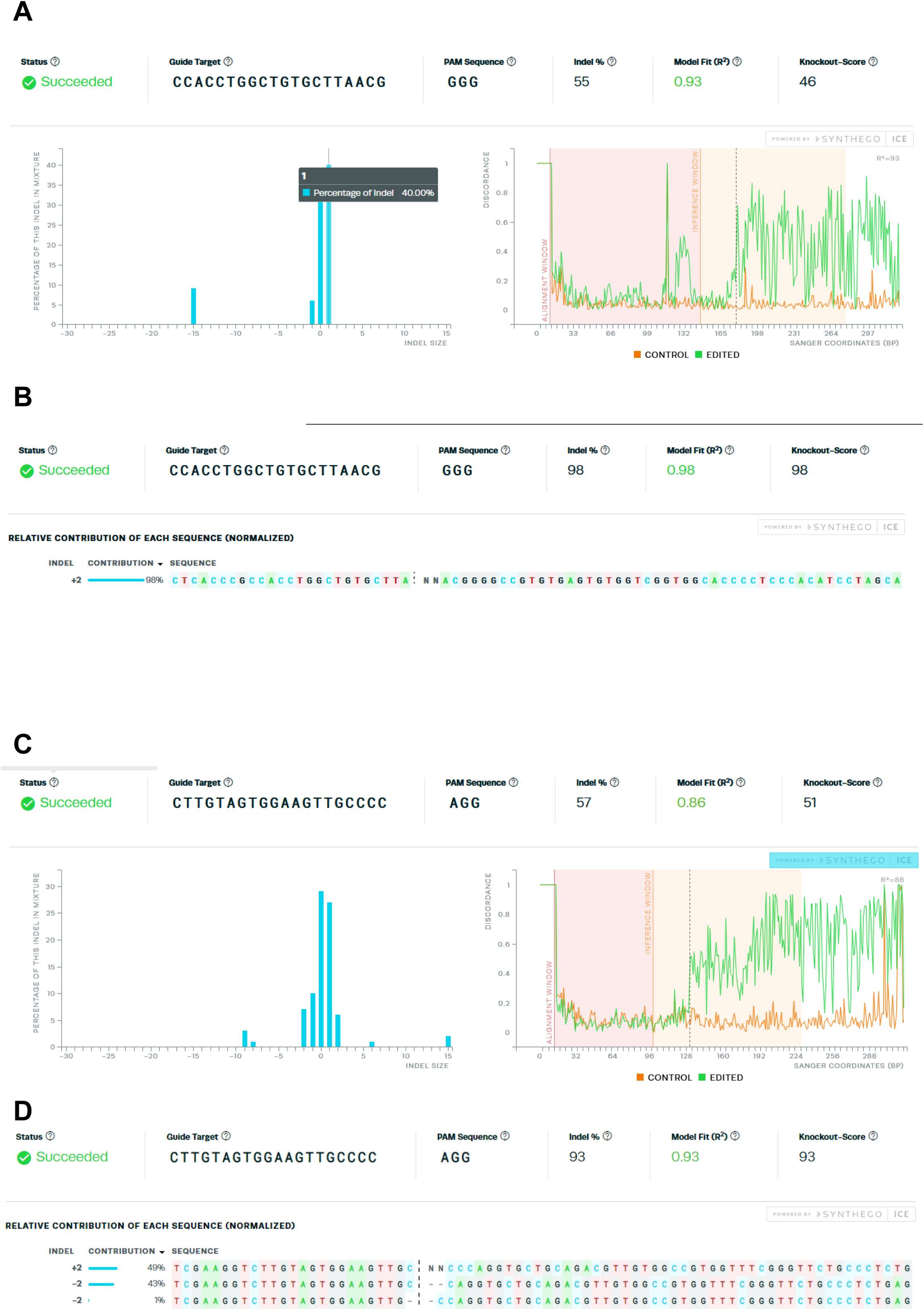
A. Ice analysis results on bulk HT-29 using gRNA2. B. Ice analysis results of HT-29 clone 22 using gRNA2. C. Ice analysis results on bulk LS-174T using gRNA3. B. Ice analysis results of HT-29 clone 11 using gRNA3.

**Supplementary Figure 6.**
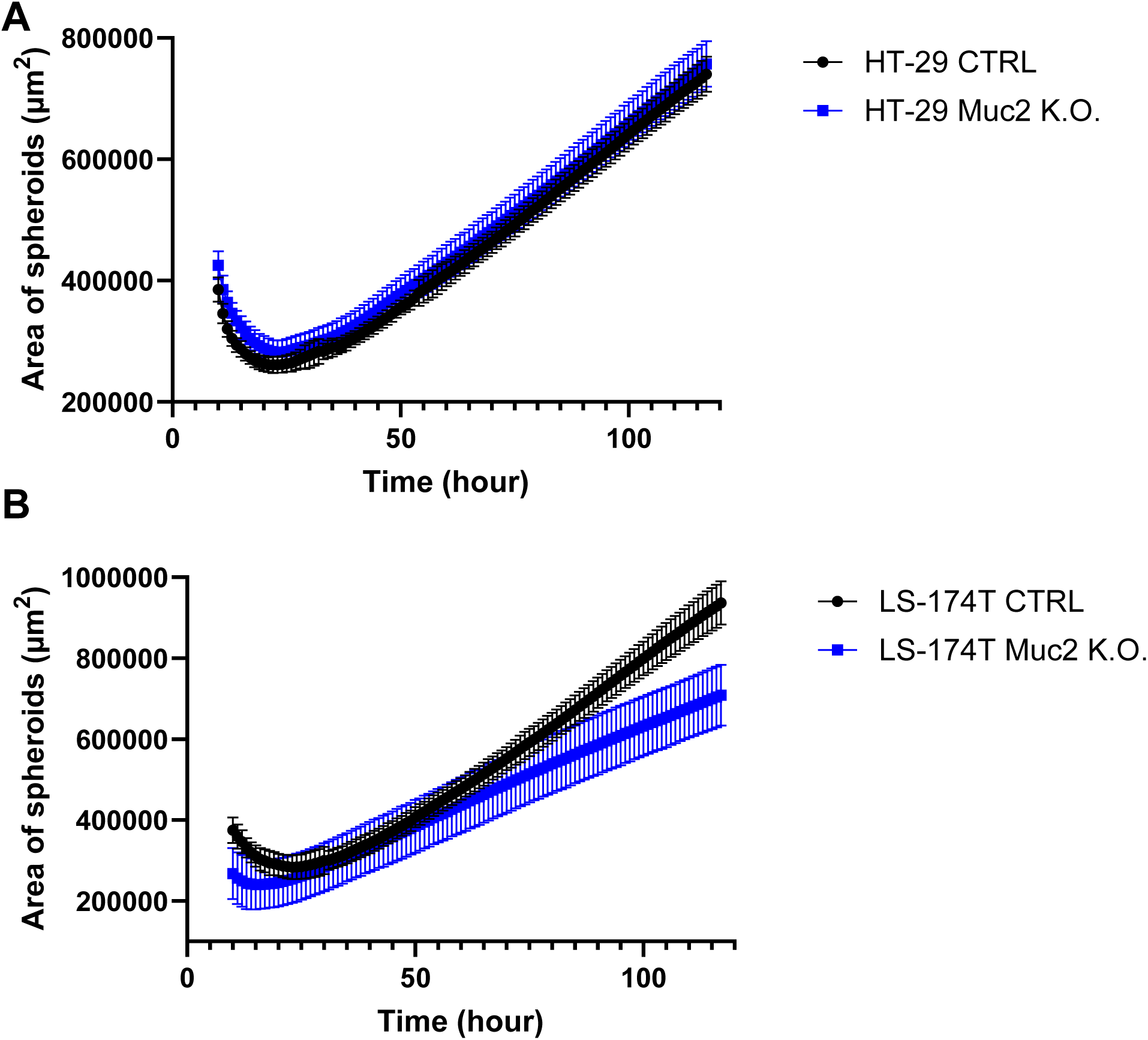
A. Surface area analysis of HT-29 CTRL and HT-29 MUC2 K.O. spheroids on day 5 (N=12 biological replicates). B. Surface area analysis of LS-174T CTRL and HT-29 MUC2 K.O. spheroids on day 5 (N=12 biological replicates).

**Supplementary Figure 7.**
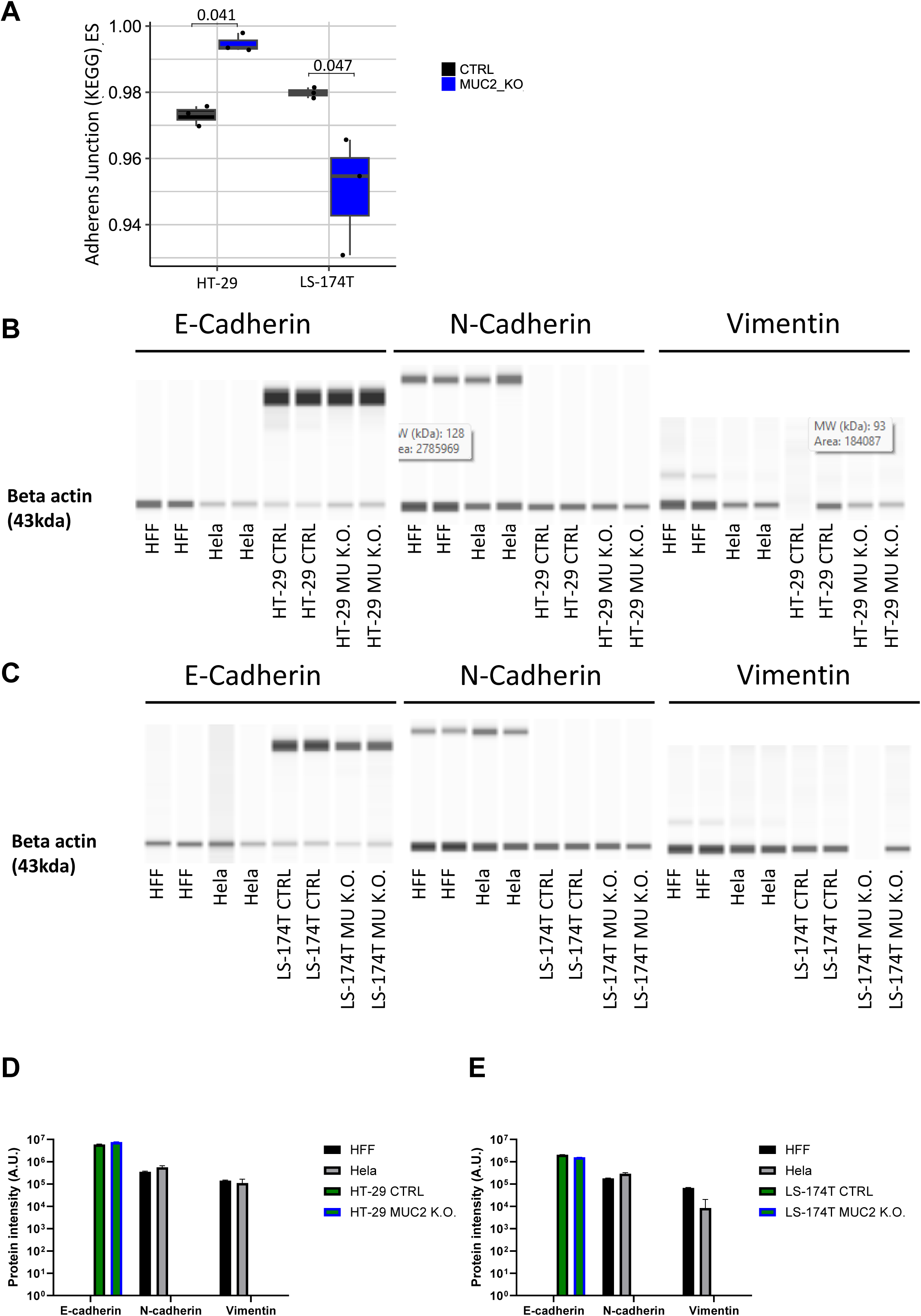
A. Box plot of adherence junction enrichment score (ES) in HT-29 and LS-174T. The ES is up-regulated in HT-29 MUC2 K.O. compared to CTRL (p=0.041) (student t-test). In contrast, the ES is down-regulated in LS-174T MUC2 K.O. compared to CTRL (p=0.047) (student t-test). B. Representative capillary western blot (CWB) results of E-cadherin, N-cadherin and Vimentin expression in human fetal fibroblast (HFF), Hela, HT-29 CTRL and HT-29 MUC2 K.O. (N=2 technical replicates). C. Representative capillary western blot (CWB) results of E-cadherin, N-cadherin and Vimentin expression in HFF, Hela, LS-174T CTRL and LS-174T MUC2 K.O. (N=2 technical replicates). D. Quantification of band intensity of E-cadherin, N-cadherin and Vimentin normalized to beta actin expression in HFF, Hela, HT-29 CTRL and HT-29 MUC2. (N=2, technical replicates). E. Quantification of band intensity of E-cadherin, N-cadherin and Vimentin normalized to beta actin expression in HFF, Hela, LS-174T CTRL and LS-174T MUC2 K.O.. (N=2, technical replicates).

**Supplementary Figure 8.**
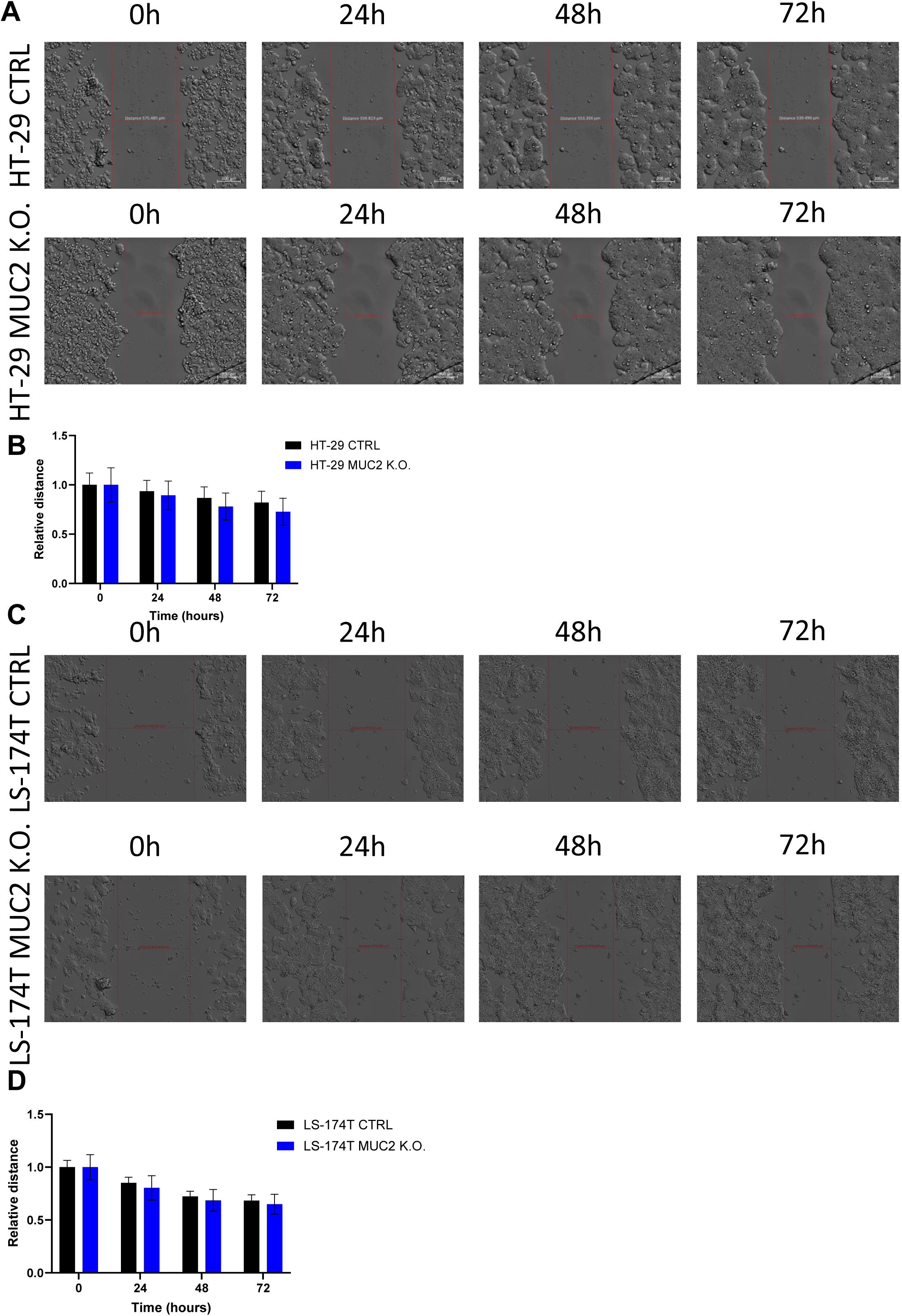
A. Representative micrographs of one of the individual experiments at time points 0, 24, 48 and 72 h in HT-29 CTRL and HT-29 MUC2 K.O. No cell motility toward the gap area was seen even after 72h, despite a clear cell multiplication. B. Size of scratch in HT-29 CTRL and MUC2 K.O. relative to 0 hour (N=12). C. Representative micrographs of one of the individual experiments at time points 0, 24, 48 and 72 h in LS-174T CTRL and LS-174T MUC2 K.O. No cell motility toward the gap area was seen even after 72h, despite a clear cell multiplication. D. Size of scratch in LS-174T CTRL and MUC2 K.O. relative to 0 hour (N=16).

**Supplementary Figure 9.**
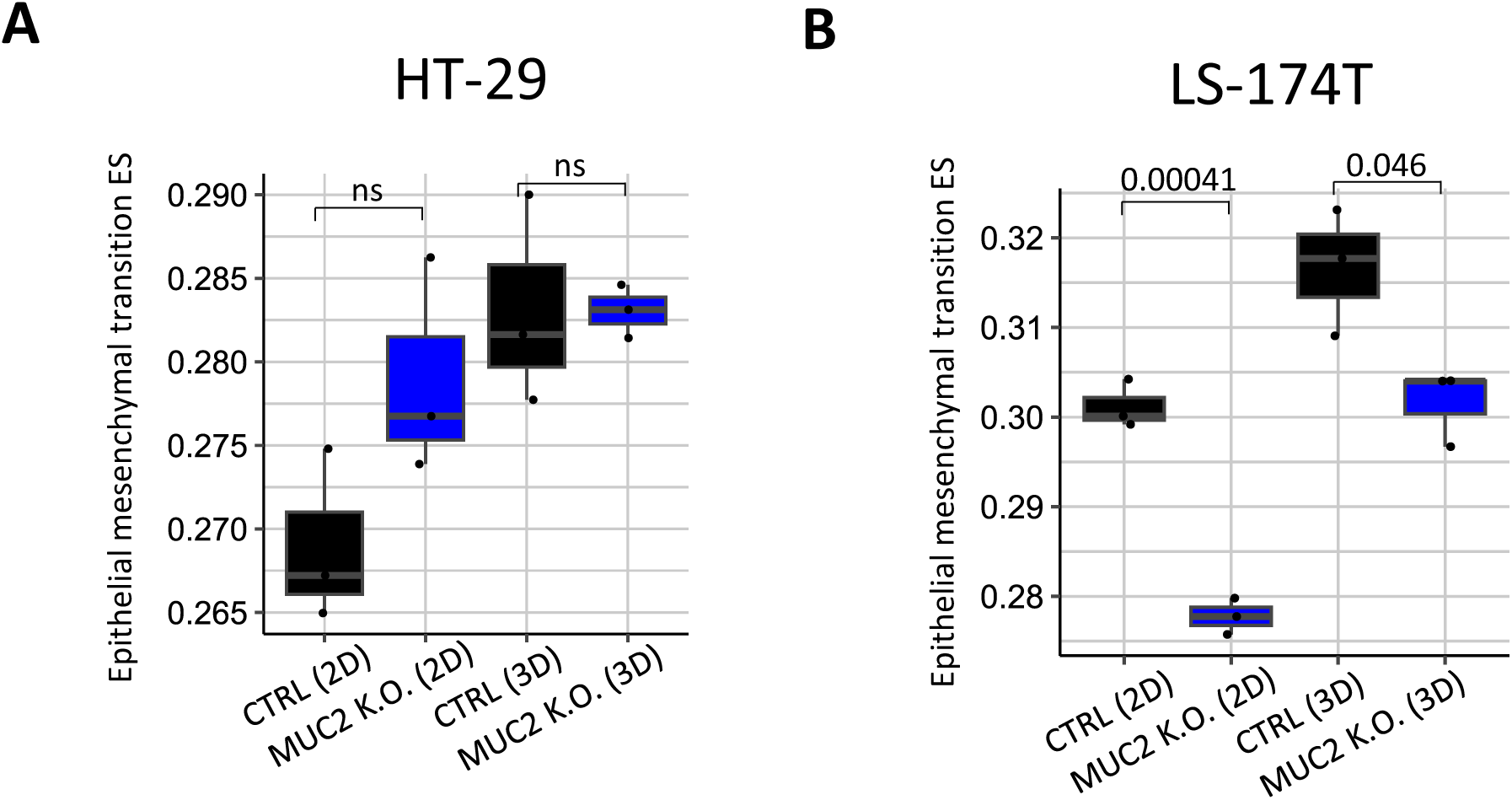
Comparison of epithelial mesenchymal transition enrichment score between CTRL (2D) and MUC2 K.O. (2D), and between CTRL (3D) and MUC2 K.O. (3D) in A. HT-29, and B. LS-174T. P-value from student t-test.

**Supplementary Figure 10.**
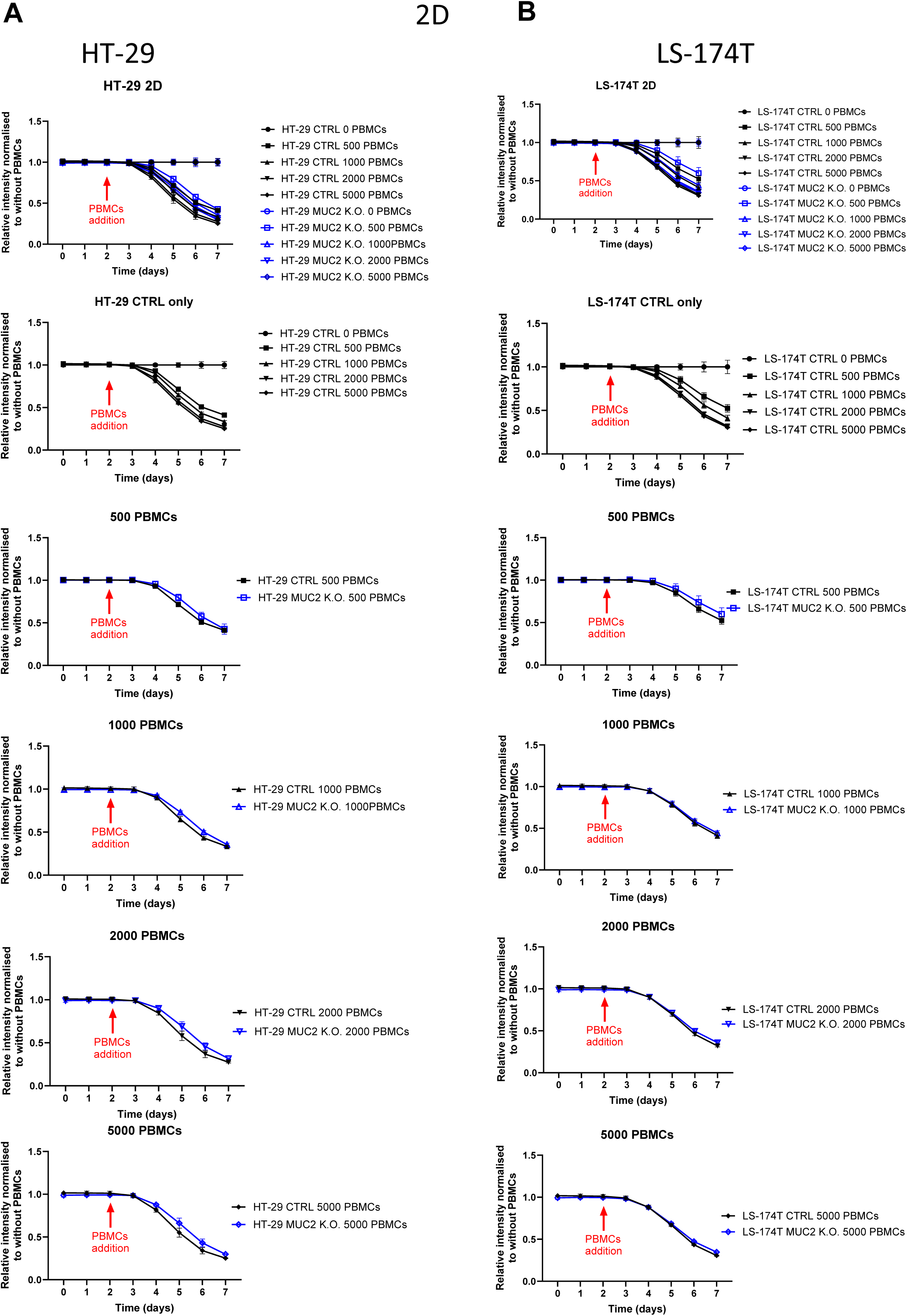
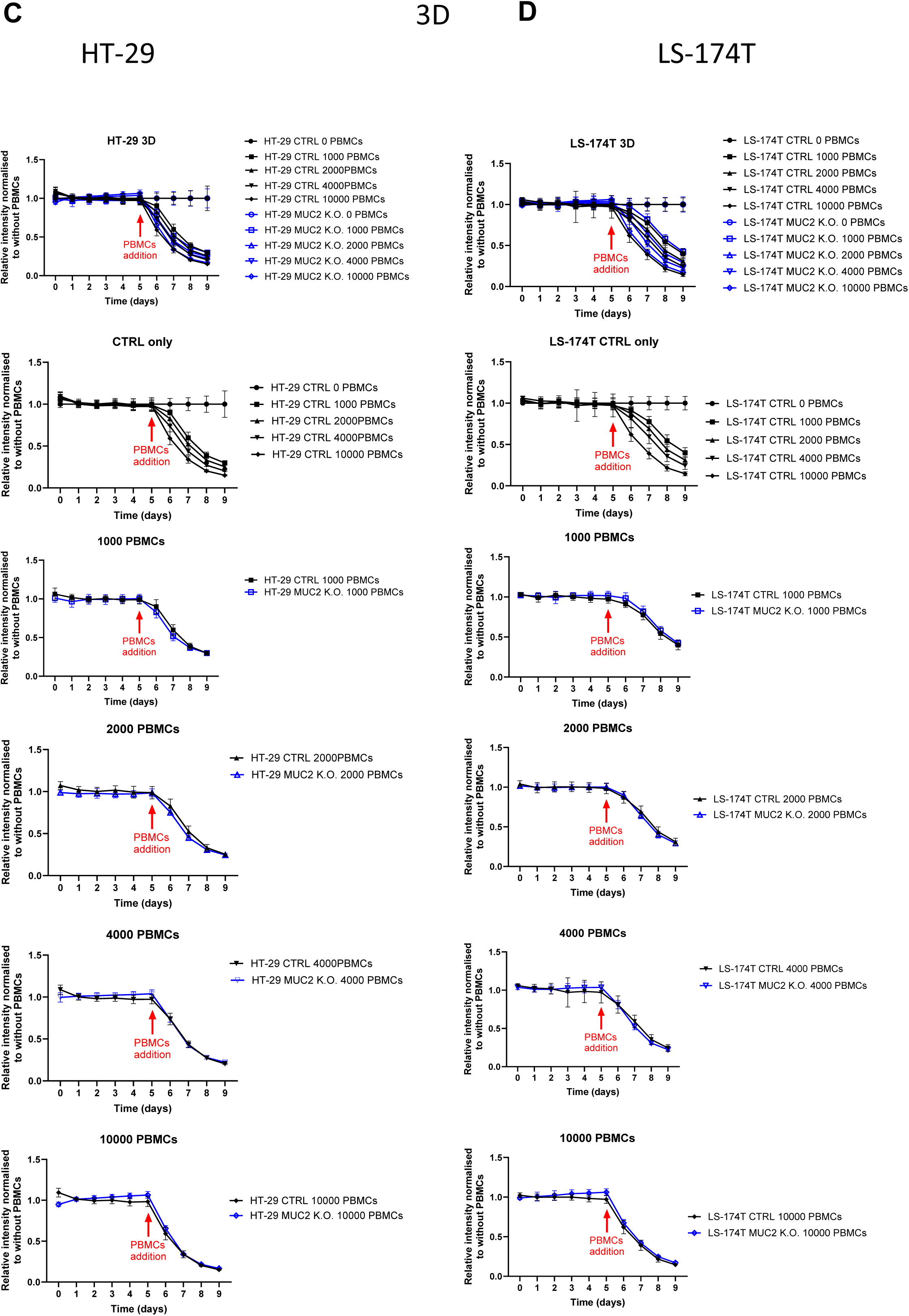
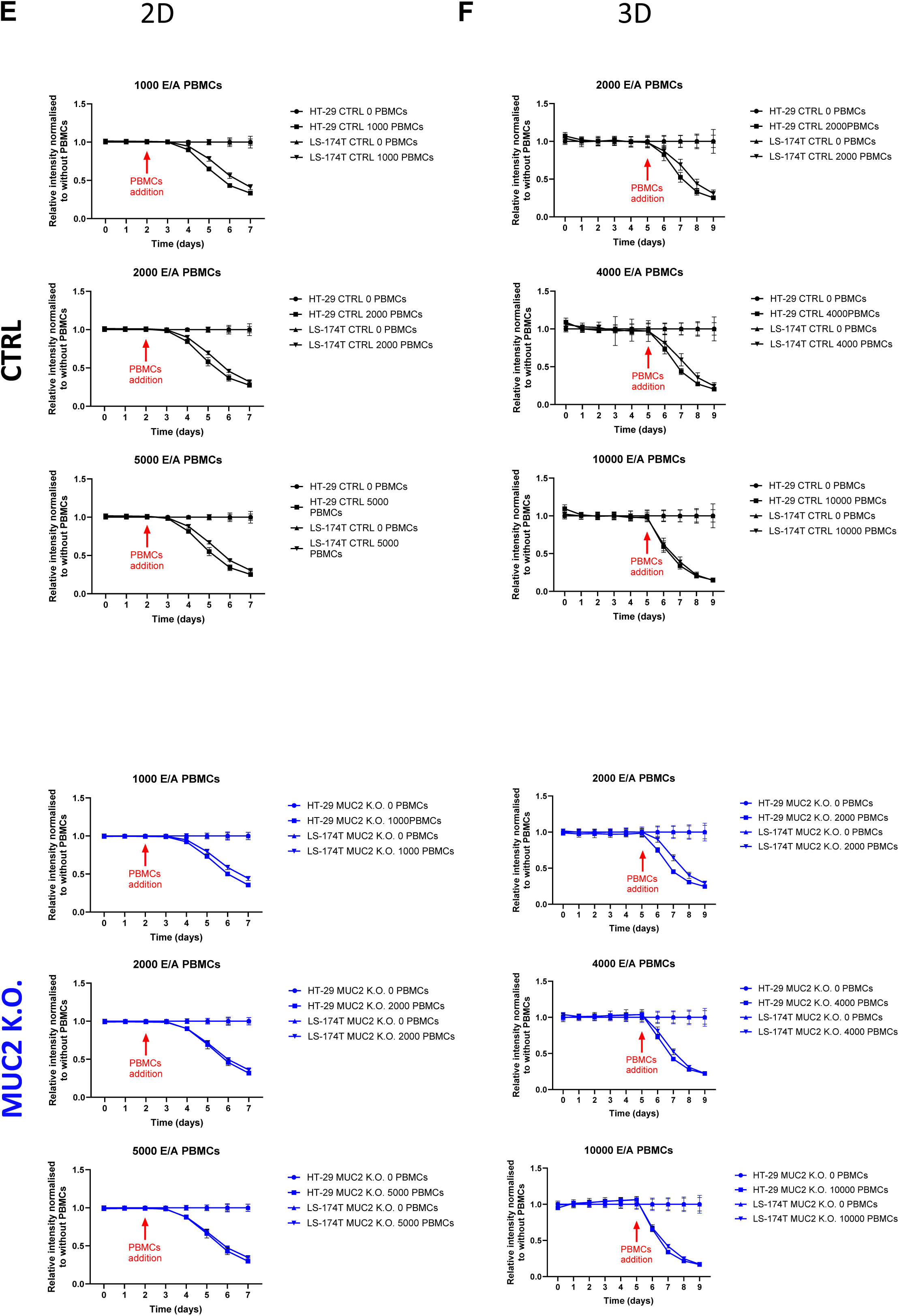
A. HT-29 CTRL and MUC2 K.O. growth and death monitored by GFP expression over 7 days in 2D set up. Top panel shows the GFP reading for HT-29 CTRL only with 0 to 10.000 E/A PBMCs added on day 2 the following panel represents the comparison of HT-29 CTRL and HT-29 MUC2 K.O. with a defined number of E/A PBCMs. Data are presented as relative to the same cell line without co-culture with PBMCs (N=18, 3 biological replicates of 6 technical replicates). B. LS-174T CTRL and MUC2 K.O. growth and death monitored by GFP expression over 7 days in 2D set up. Top panel shows the GFP reading for LS-174T CTRL only with 0 to 10.000 E/A PBMCs added on day 2 the following panel represents the comparison of LS-174T CTRL and LS-174T MUC2 K.O. with a defined number of E/A PBCMs. Data are presented as relative to the same cell line without co-culture with PBMCs (N=18, 3 biological replicates of 6 technical replicates). C. HT-29 CTRL and MUC2 K.O. growth and death monitored by GFP expression over 9 days in 3D set up. Top panel shows the GFP reading for HT-29 CTRL only with 0 to 10.000 E/A PBMCs added on day 5 the following panel represents the comparison of HT-29 CTRL and HT-29 MUC2 K.O. with a defined number of E/A PBCMs. Data are presented as relative to the same cell line without co-culture with PBMCs (N=18, 3 biological replicates of 6 technical replicates). D. LS-174T CTRL and MUC2 K.O. growth and death monitored by GFP expression over 7 days in 3D set up. Top panel shows the GFP reading for LS-174T CTRL only with 0 to 10.000 E/A PBMCs added on day 5 the following panel represents the comparison of LS-174T CTRL and LS-174T MUC2 K.O. with a defined number of E/A PBCMs. Data are presented as relative to the same cell line without co-culture with PBMCs (N=18, 3 biological replicates of 6 technical replicates). E. Comparison of HT-29 with LS-174T CTRL or MUC2 K.O. with various number of E/A PBMCs in 2D. Data are presented as relative to the same cell line without co-culture with PBMCs (0 E/A PBMCs) (N=18, 3 biological replicates of 6 technical replicates). F. Comparison of HT-29 with LS-174T CTRL or MUC2 K.O. with various number of E/A PBMCs in 3D. Data are presented as relative to the same cell line without co-culture with PBMCs (0 E/A PBMCs) (N=18, 3 biological replicates of 6 technical replicates).

**Supplementary Figure 11.**
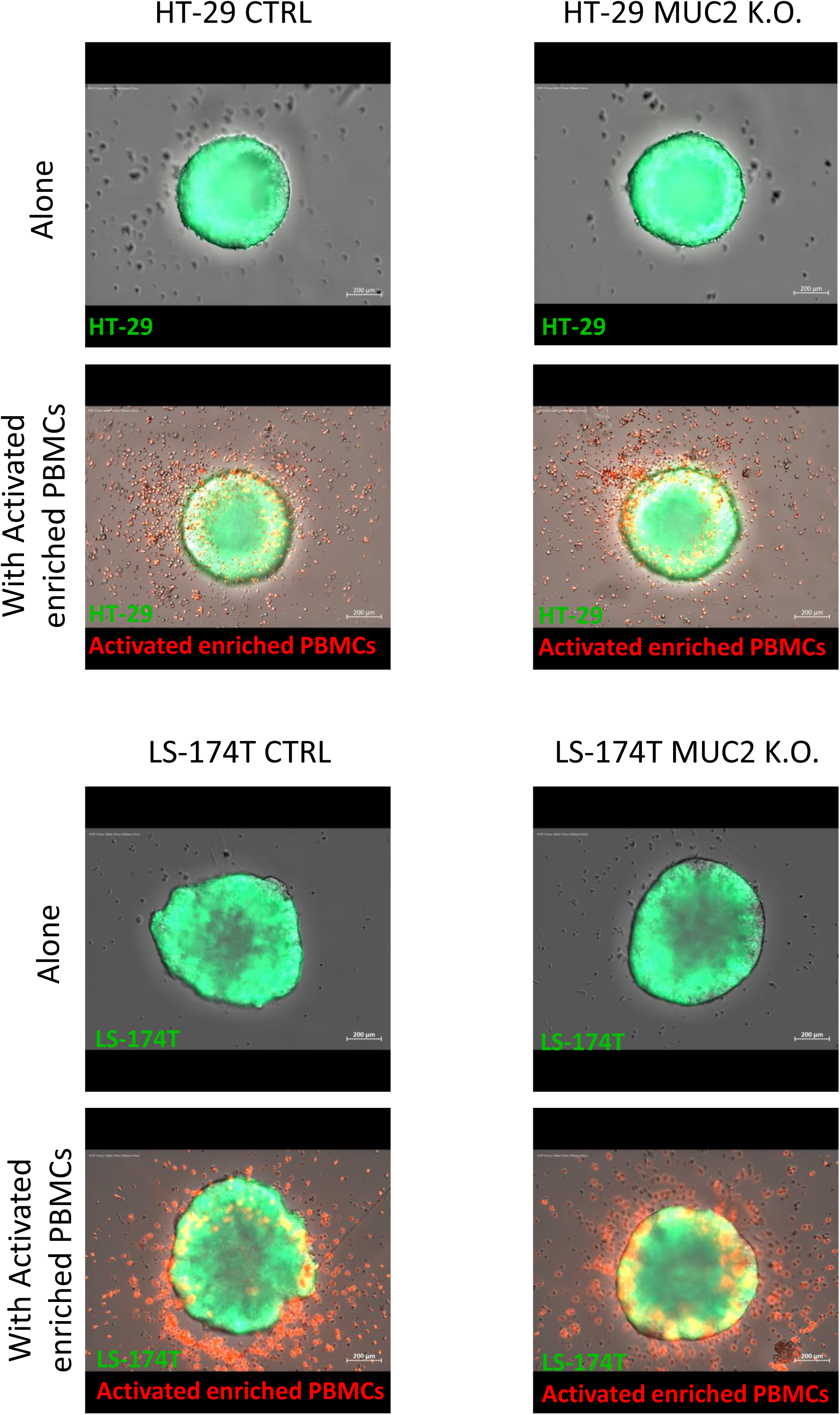
Representative video of HT-29 CTRL, HT-29 MUC2 K.O., LS-174T CTRL, LS-174T MUC2 K.O. from day 5 to day 7 with and without E/A PBMCs.

**Supplementary Figure 12.**
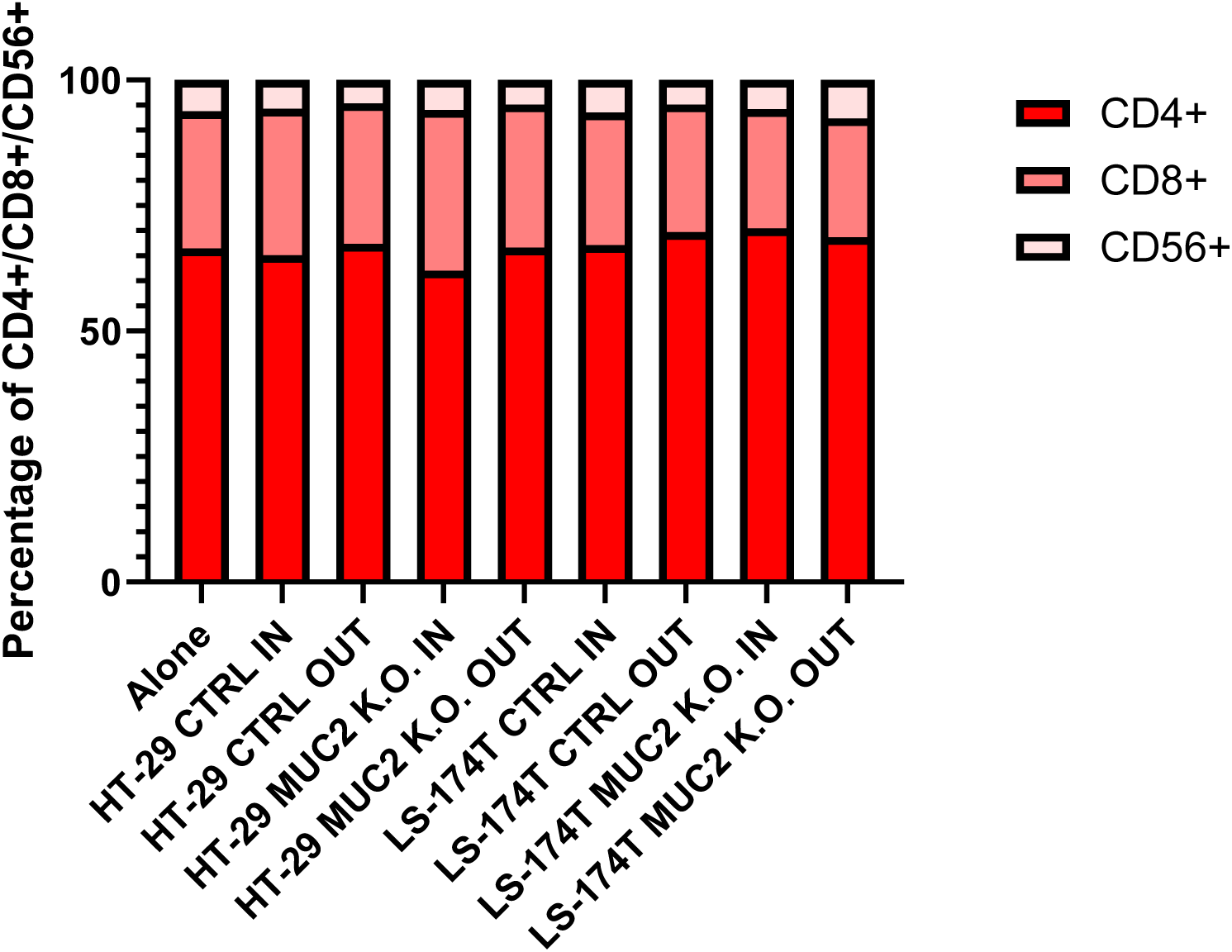
Relative proportion in percentage of CD4+, Cd8+ and CD56+ cells among the CD45+ cells in the IN and OUT fraction after co-culture with HT-29 and LS-174T CTRL or MUC2 K.O. we see no enrichment of specific selection of either cell type in the IN or OUT fraction compared to E/A PBMCs alone.

**Supplementary Figure 13.**
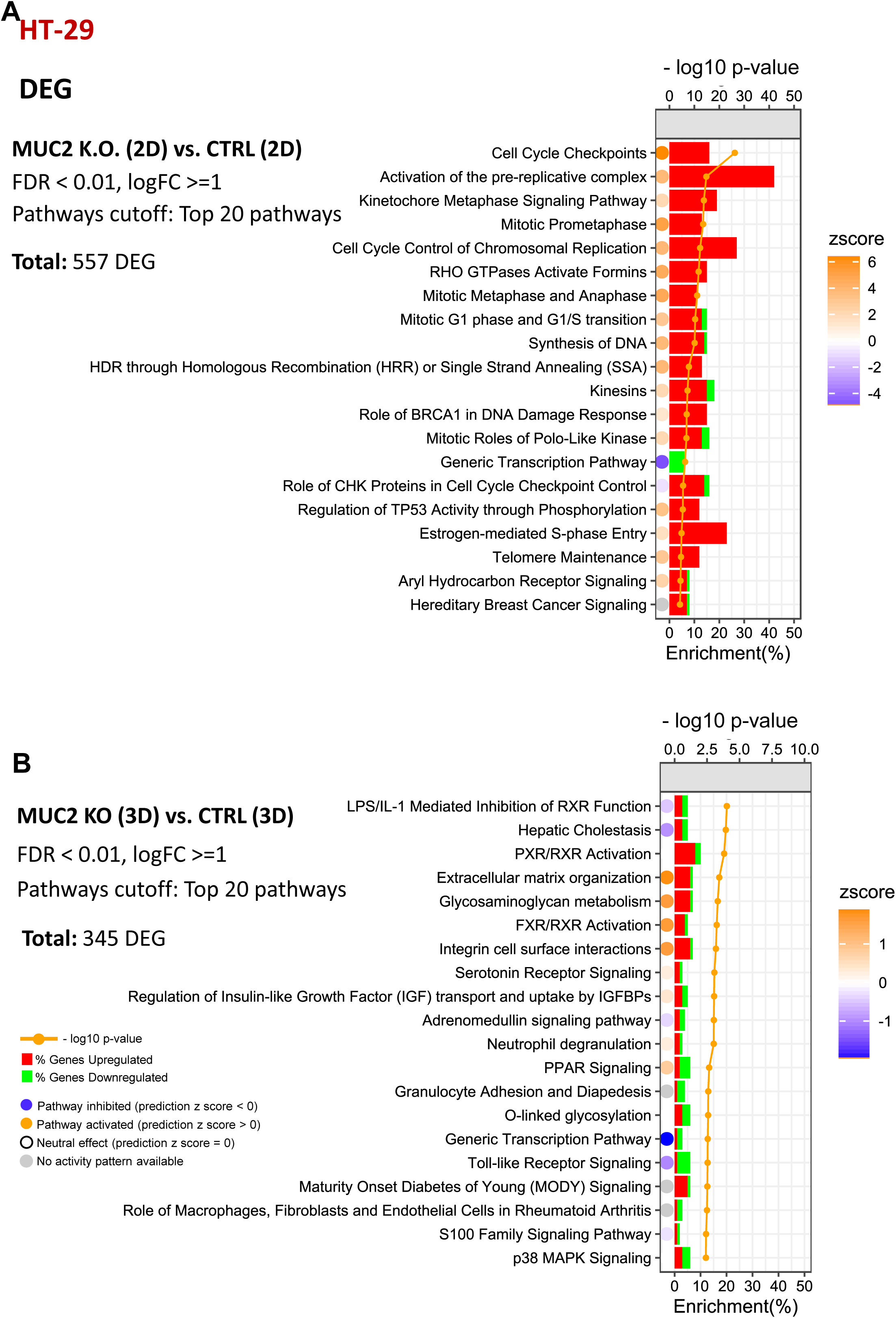
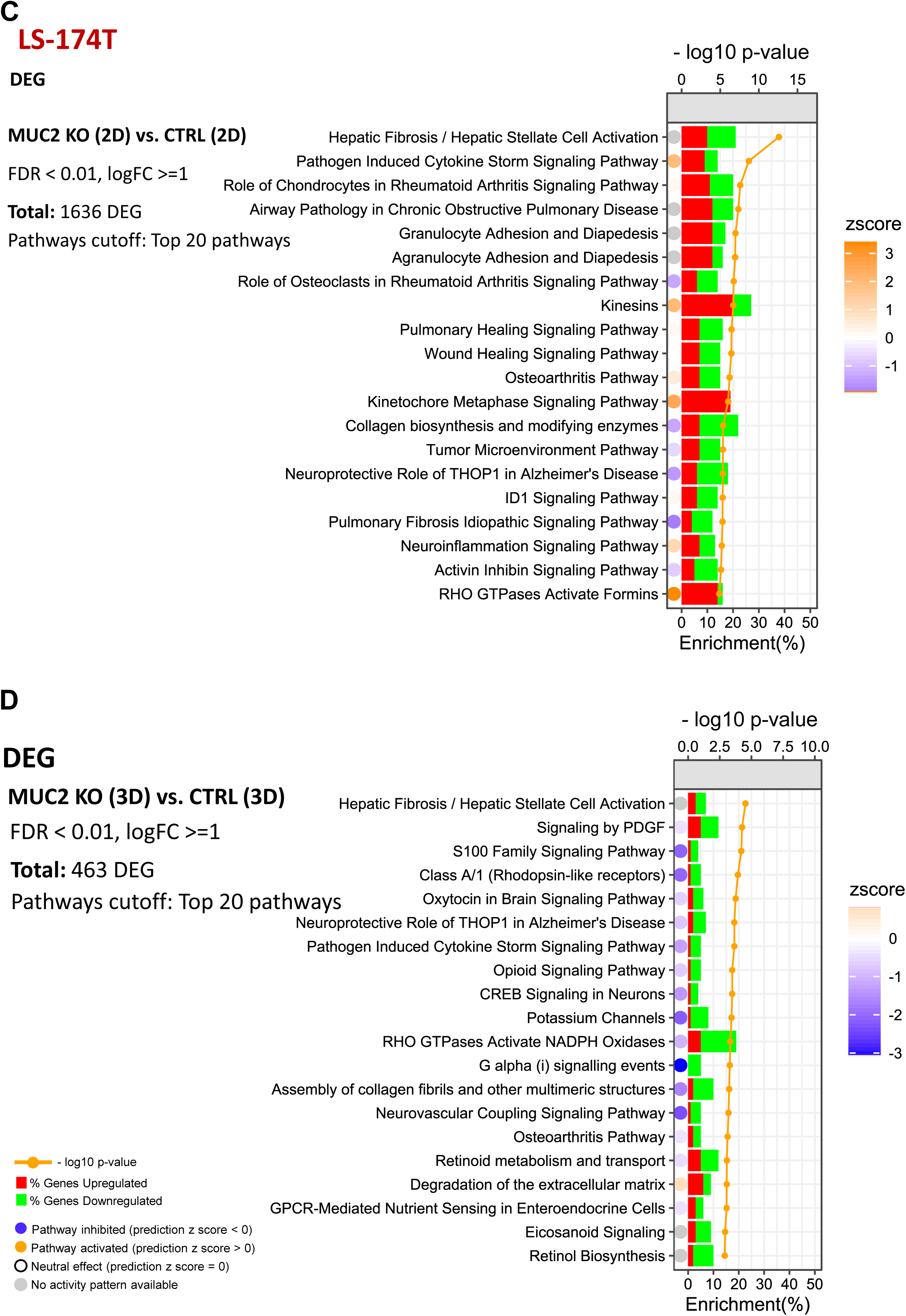
IPA pathways enrichment analysis. Top 20 significant pathways based on p-value associated with DEG (FDR < 0.01, logFC >=1) between MUC2 K.O. (2D) and CTRL (2D) in A. HT-29 (n = 557) C. LS-174T (n = 1636). Between MUC2 K.O (3D) and CTRL (3D) in B. HT-29 (n = 345) D. LS-174T (n = 463). Histograms represent the proportion (%) of DEGs upregulated (red) or downregulated (green) in MUC2 K.O versus CTRL. The circles represent the pathway activation status. Blue circle indicates the pathway is inhibited with a negative z-score, orange circle represents a pathway is activated with a positive z-score, the white circle represents the pathway is neutral with zero z-score, while a gray circle indicates that the pathway activity is unknown.

**Supplementary Figure 14.**
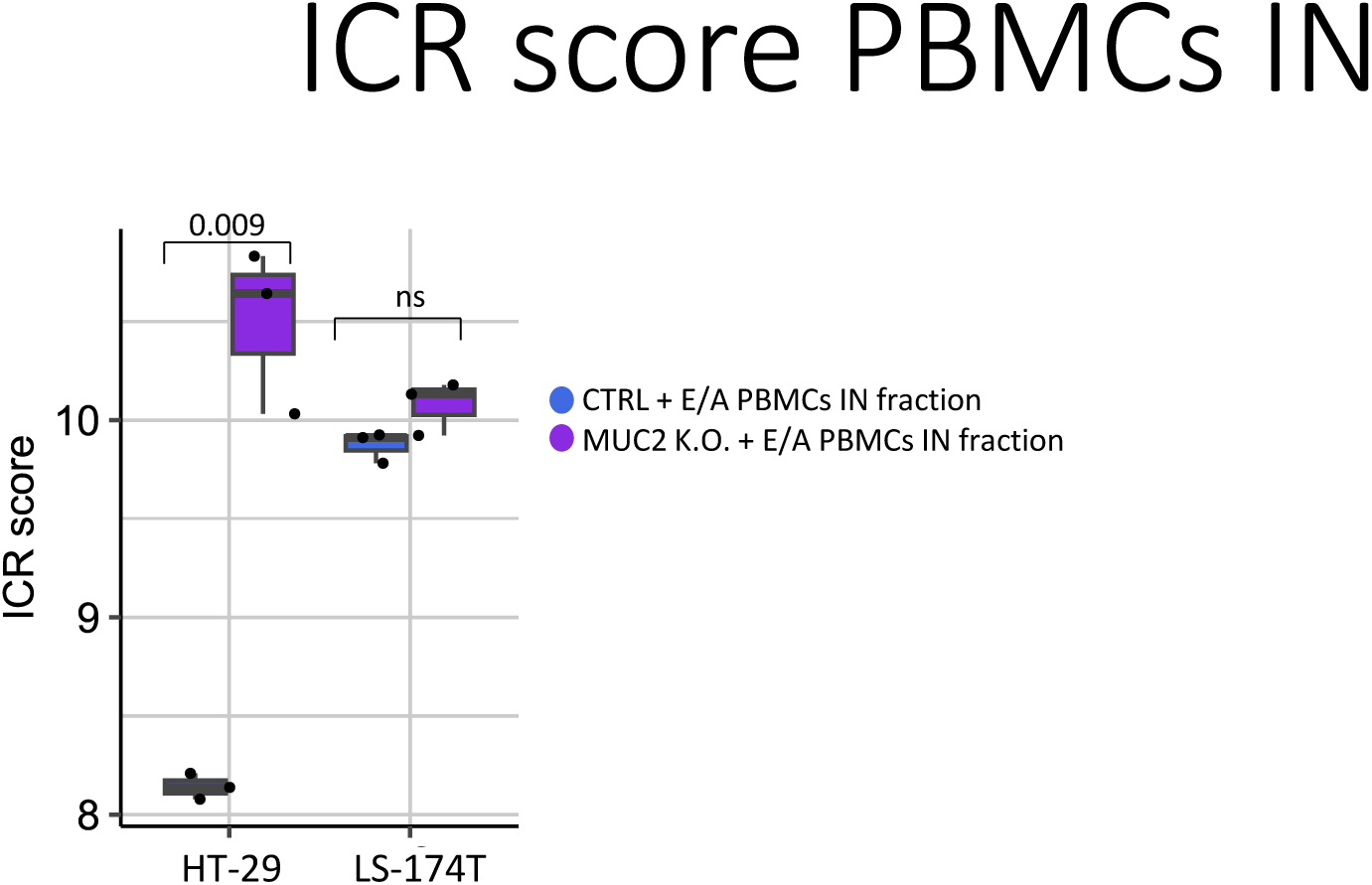
Comparison of ICR score mean value between CTRL + E/A PBMCs IN fraction and MUC2 K.O. + E/A PBMCs IN fraction in HT-29 and LS-174T. P-value from student t-test.

**Supplementary Figure 15.**
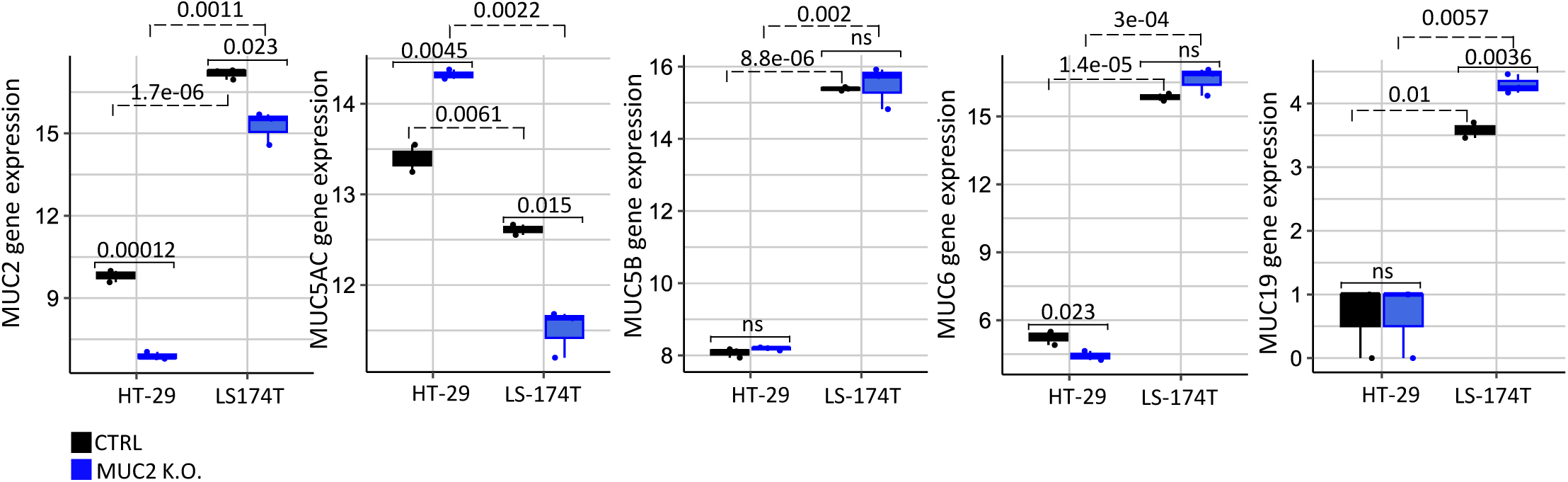
Comparison of genes expression of other gel forming mucin in HT-29 and LS-174T spheroids between MUC2 K.O. (in blue) and CTRL (in black) (3D). Statistical analysis represented as p-values from student’s t-test.

## Ethics approval and consent to participate

“Not applicable”

## Consent for publication

“Not applicable”

## Availability of data and materials

All data produced in the present study are available upon reasonable request to the authors.

## Acknowledgements

The authors would like to acknowledge the Sidra Medicine research branch core facilities, especially the genomic core and our research administration team without whom we could not have performed this work.

## Author’s contribution

C.R., A.J., S.H., A.S. performed the experiments, C.R., A.J., E.A., S.S., performed the analysis C.R., A.J., E.A. drafted the manuscript and the figures, W.H., edited the text. A.A, B.L. revised the manuscript and provided input. Conceptualization and project supervision by D.B., C.R., and W.H., project coordination C.R. All authors reviewed the manuscript.

